# Do Newly Born Orphan Proteins Resemble Never Born Proteins? A Study Using Three Deep Learning Algorithms

**DOI:** 10.1101/2022.08.02.502493

**Authors:** Jing Liu, Rongqing Yuan, Wei Shao, Jitong Wang, Israel Silman, Joel L. Sussman

## Abstract

*‘Newly Born’* proteins, devoid of detectable homology to any other proteins, known as orphan proteins, occur in a single species or within a taxonomically restricted gene family. They are generated by expression of novel Open Reading Frames, and appear throughout evolution. We were curious if the three recently developed programs for predicting protein structures, viz., AlphaFold2, RoseTTAFold, and ESMFold, might be of value for comparison of such ‘*Newly Born’* proteins to random polypeptides with amino acid content similar to that of native proteins, which have been called ‘*Never Born*’ proteins. The programs were used to compare the structures of two sets of ‘*Never Born’* proteins that had been expressed – Group 1, which had been shown experimentally to possess substantial secondary structure, and Group 3, which had been shown to be intrinsically disordered. Overall, the models generated were scored as being of low quality but revealed some general principles. Specifically, all four members of Group 1 were predicted to be compact by all three algorithms. The members of Group 3 were predicted to be very extended, as would be expected for intrinsically disordered proteins. The three programs were then used to predict the structures of three orphan proteins whose crystal structures had been solved, two of which display novel folds. Finally, they were used to predict the structures of seven orphan proteins with well-identified biological functions, whose 3D structures are not known. Two proteins, which were predicted to be disordered based on their sequences, are predicted by all three structure algorithms to be extended structures. The other five were predicted to be compact structures with two exceptions in the case of AlphaFold2. All three prediction algorithms make remarkably similar and high-quality predictions for one large protein, HCO_11565, from a nematode. It is conjectured that this is due to many homologs in the taxonomically restricted family of which it is a member and to the fact that the *Dali* server revealed several non-related proteins with similar folds. Overall, orphan and taxonomically restricted proteins are often predicted to have compact 3D structures, sometimes with a novel fold that is a consequence of their novel sequences, which are associated with the appearance of new biological functions.

## 1. INTRODUCTION

The accepted view, until quite recently, has been that protein sequences have evolved so as to incorporate the features required for optimal folding and function^1^. Specific amino acid or oligopeptide patterns appear to yield insights into phylogenetic differences between the three kingdoms: prokaryotes, archaea, and eukaryotes^2^. Surprisingly, however, evidence has been presented that ‘from a sequence similarity perspective, *real unrelated* proteins are indistinguishable from random amino acid sequences’^3^. This, at first sight, seems counterintuitive, because it might be anticipated that natural sequences would differ from random sequences in their folding characteristics. Indeed, this conclusion has been challenged by using an *ad hoc* Evolutionary Neural Network Algorithm (ENNA) to assess whether and to what extent natural proteins can be distinguished from random sequences^4^. The ENNA approach could correctly distinguish natural proteins from random sequences with >94% accuracy. Very recently, a distilled protein language model was used to distinguish natural from random sequences with >92% accuracy^5^.

Two sets of studies have shown that random sequences with native**-**like amino acid composition can be expressed and that in many cases, the expressed polypeptide chains of these *‘Never Born’* proteins^6^ fold in aqueous solution into compact structures that display resistance to proteolysis^7^, or have substantial secondary structure elements^8^. For some earlier studies using randomized sequences see^9-11^.

Recently, 2,000 random sequences of 100 amino acids each were used to generate 3D models with RoseTTAFold^12^. These initial models were optimized by Monte Carlo sampling in amino acid sequence space to yield “novel proteins spanning a wide range of sequences and predicted structures”^13^. 129 of these sequences were then expressed in *E. coli*. Of those expressed, 27 yielded monodisperse species with circular dichroism spectra consistent with a native structure. Three of the 3D structures of three were determined, and all three displayed novel folds. The folding propensity of random sequences is of relevance in the context of the issue of orphan genes and of the proteins that they express, *viz*., *‘Newly Born’* proteins, devoid of detectable homology to any other proteins, which occur in a single species, or proteins that occur within a taxonomically restricted gene (TRG) family, TRGPs^14-20^. Such a possibility was considered to be impossible even by such a distinguished figure as François Jacob^21^. However, to quote a recent review - “The origin of novel protein-coding genes was once considered so improbable as to be impossible. In the last decade, and especially in the last five years, this view has been overturned by extensive evidence from diverse eukaryotic lineages”^17^. Both the term orphan gene and the term orphan protein are often loosely used in more than one context. In the present study, the term orphan gene refers to a gene for which evidence has been presented that it has arisen from what was previously a non-coding DNA sequence, and is expressed as an open reading frame (ORF). Thus, the orphan protein for which it codes is seen only in a single species or in one that is closely related taxonomically, *i*.*e*., a protein generated by a TRG, a TRGP. The general contention is thus that new genes may appear out of previously non-coding genomic regions, a process also known as exonization^22^, and code for novel protein sequences^23^. The question that then arises is how new functional protein domains might evolve out of such random sequences^14^? It is fair to say that this is still an open question and that much more experimental data will be required. Of particular interest are studies of higher primates in which novel genes were identified that are shared by chimpanzees, gorillas, and humans, whereas other *de novo* genes may, for example, be restricted to humans^24-29^. It should also be mentioned that it has been suggested that novel protein sequences may also be generated by ORFs present in long non-coding RNAs (lncRNAs)^15,30^.

Very recently, the field of structural biology has undergone a revolution due to the development of the deep-learning-based protein structure prediction programs, AlphaFold2 (AF2)^31^, RoseTTAFold (RTF)^12^ and, most recently, Evolutionary Scale Modeling (ESM-2)^32^. All three algorithms have been shown to predict 3D structures for many natural sequences closely resembling the experimental structures deposited in the PDB^33-35^. We were curious as to whether these powerful new protein structure prediction tools might be valuable for distinguishing natural from random sequences. Here, we use all three AI/Deep Learning programs just mentioned to predict the structures of several natural sequences and of ‘*Newly Born*’ orphan proteins. We also used them to predict the structures of the random sequences of ‘*Never Born*’ proteins expressed by Tretyachenko *et al*.^8^, some of which these authors had shown to fold into compact structures, others to belong to the category of intrinsically disordered proteins (IDPs)^36^.

## 2. METHODS

### 2.1. Protein sequences

Protein sequences for the crystal structures were retrieved from the PDB (https://www.rcsb.org), for the IDPs and for the ‘*Newly Born*’ orphan proteins from UniProt (https://www.uniprot.org), and for the ten ‘*Never Born*’ proteins from the supplementary information associated with the study of Tretyachenko *et al*.^8^ (https://static-content.springer.com/esm/art%3A10.1038%2Fs41598-017-15635-8/MediaObjects/41598_2017_15635_MOESM1_ESM.pdf). All these sequences are listed in Table 1.

**Table 1.**
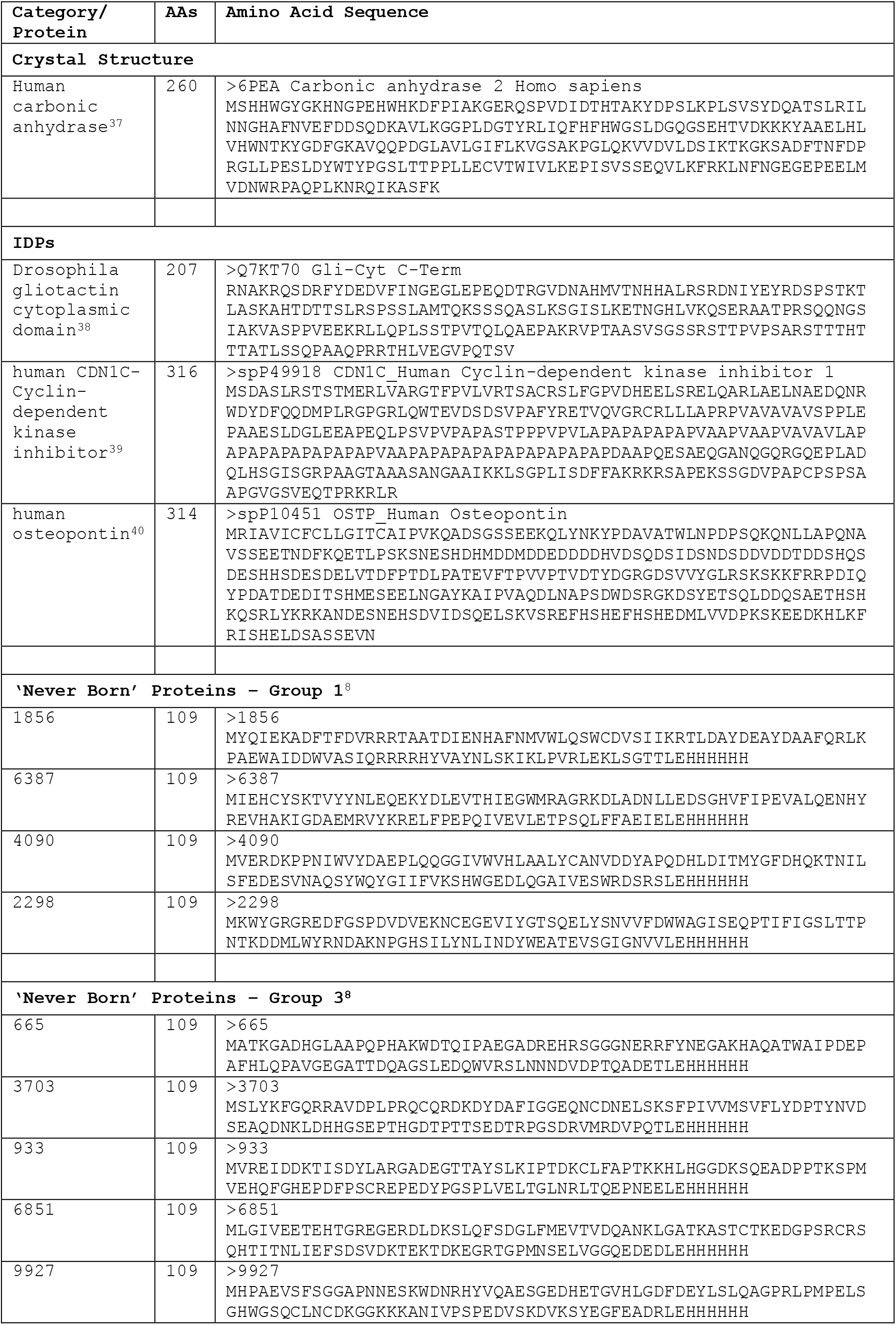
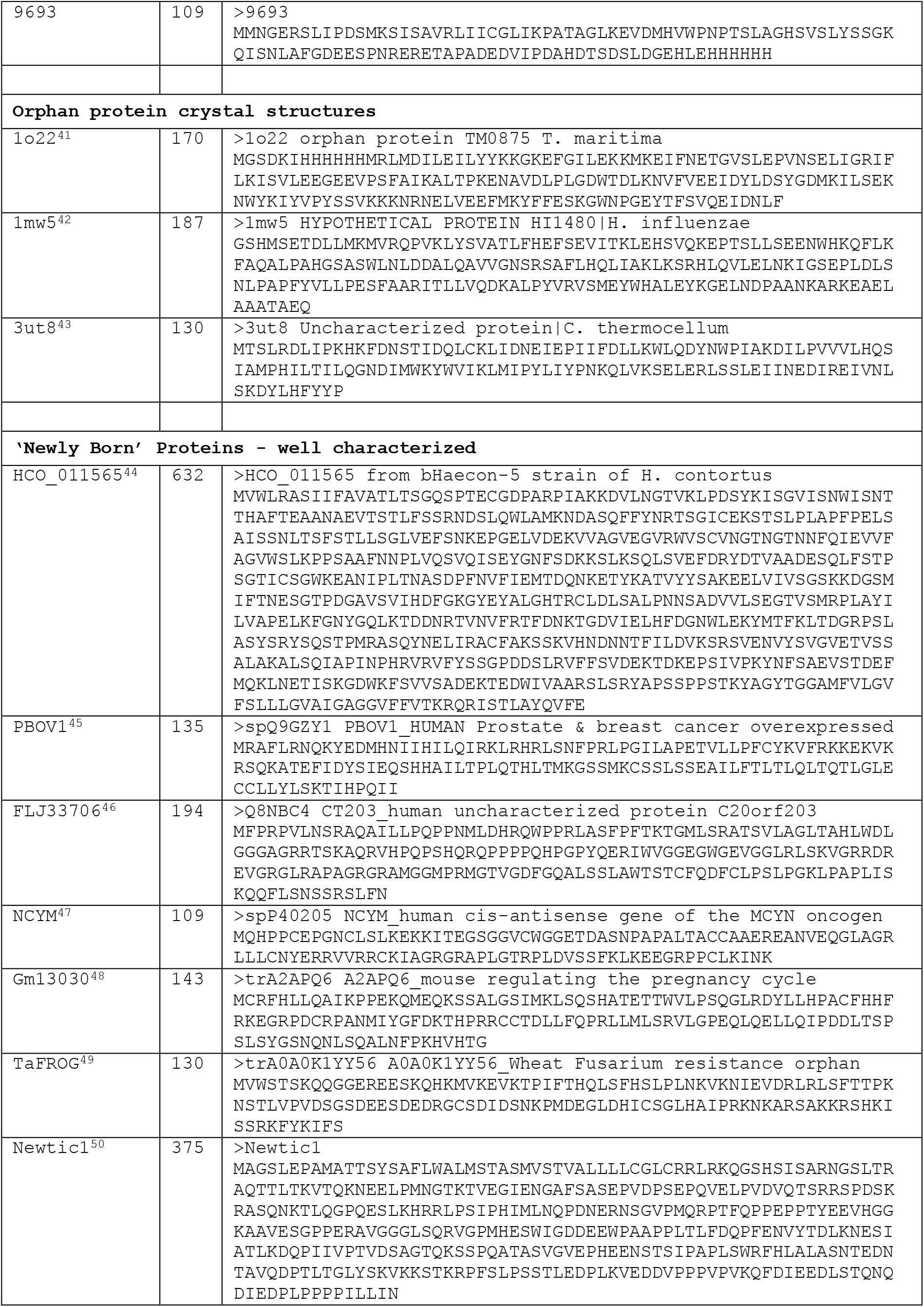
Sequences of proteins and polypeptides utilized

### 2.2. AlphaFold2 predictions

The AlphaFold2 (AF2) predictions were performed by the AlphaFold2_advanced Colab^51^ at https://colab.research.google.com/github/sokrypton/ColabFold/blob/main/beta/AlphaFold2_advanced.ipynb. The defaults used were:

- multisequence alignment, mmseq2
- template protein structures were not used.

### 2.3. Evolutionary scale modeling

The Evolutionary Scale Modeling (ESM-2) predictions were performed via the *esmfold*.*py* plug-in to PyMp; [https://github.com/JinyuanSun/PymolFold] for sequences containing up to 400 amino acids, and for longer sequences by ESMFold.ipynb Colab [https://colab.research.google.com/github/sokrypton/ColabFold/blob/main/ESMFold.ipynb]. The defaults for this server were employed.

### 2.4. RoseTTAFold predictions

The RoseTTAFold (RTF) predictions were performed by the Robetta server using the RoseTTAFold option^12^ at: https://robetta.bakerlab.org. The defaults for this server were employed.

### 2.5. Natural protein sequence

As a control, we selected for this study one native globular protein whose crystal structure has been experimentally determined to high resolution, 1.36Å, human carbonic anhydrase (UniProt - P00918, PDB - 6pea). The sequence information for this protein was obtained from the UniProt database (https://www.uniprot.org) and the crystal structure from the PDB [https://www.rcsb.org].

### 2.6. Structure prediction and comparison

For each of the sequences submitted to the AF2 and RTF servers, the 5 most probable models were generated, while ESM-2 generated only a single model. In the cases in which the 3D structures were available, the predicted structures were aligned with the experimental structures using PyMol [PyMOL Molecular Graphics System, Version 2.1 ATI-4.8.101, Schrödinger, LLC].

AF2 and ESM-2 produce an estimate of the confidence, on a scale of 0-100, for each residue. This confidence measure is called pLDDT, as defined on the IDDT-Cα metric^52^. It is stored in the B-factor fields of the corresponding PDB files. pLDDT is also used to color-code the residues of the model in the 3D structure viewer (Fig. 1).

**Fig. 1.**
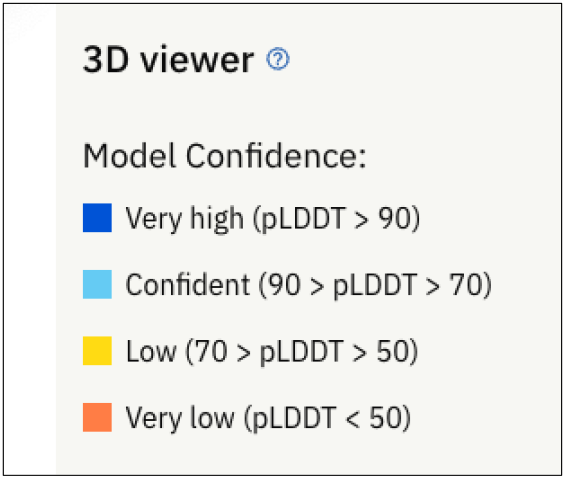
Color coding scheme based on the PDBe AlphaFold database [https://alphafold.ebi.ac.uk]

For models predicted by RTF, the RMSD values are inserted in place of B-factors in the PDB files generated. They are then converted into pLDDT values using a Python program that we wrote, based on the formulae described by^53^, as seen at https://phenix-online.org/version_docs/dev-4380/reference/process_predicted_model.html.

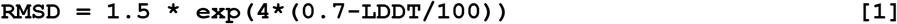

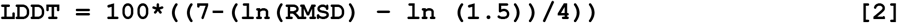

To render the coloring schemes consistent for all structures shown, the B-factors in the original PDB files for experimentally determined 3D structures were converted to pLDDT using the following equations:

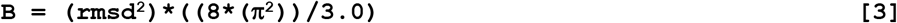

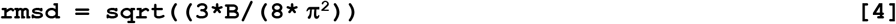

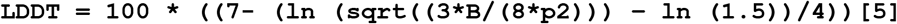

For all structures, we then applied the color scheme used in the PDBe AlphaFold2 database. [https://alphafold.ebi.ac.uk] shown in Fig. 1, via the *color_plddt*.*py* plugin for PyMol [https://github.com/JinyuanSun/PymolFold].

### 2.7. Detection of novel and unique folds

The Dali server (http://www2.ebi.ac.uk/dali) was used to check if the folds were novel^54,55^.

### 2.8. Calculation of Accessible Surface Area

The Accessible Surface Area (ASA) for 3D protein structures was calculated using the PyMOL Molecular Graphics System, Version 2.1 ATI-4.8.101 Schrödinger, LLC].

### 2.9. Morphs for the 5 top models

To facilitate comparison of each set of the 5 top models generated by AF2 and RTF, a morph was generated via PyMOL after first aligning the 5 top model structures on top of each other using PyMOL. These morphs are displayed under Supplementary Information.

### 2.10. Prediction of intrinsically disordered regions

The prediction of intrinsically disordered regions was performed using FoldIndex^56^ (https://fold.proteopedia.org/cgi-bin/findex) and flDPnn^57^ (http://biomine.cs.vcu.edu/servers/flDPnn).

### 2.11. Amino acid compositions, pIs, and charge calculations

Amino acid compositions were calculated using the Expasy ProtParam tool [https://web.expasy.org/protparam]. The pI values, and the protein charges at pH 7.4, were calculated using the Prot pi | Protein Tool [https://www.protpi.ch/Calculator/ProteinTool].

### 2.12. BLASTP sequence searches

BLASTP sequence searches were performed using the NIH-NLM site [https://blast.ncbi.nlm.nih.gov/Blast.cgi?PAGE=Proteins].

#### 3. RESULTS

The AI/Deep Learning tools that have recently developed have revolutionized the prediction of 3D protein structures^31,32,53^. This is exemplified by Fig. 2, which displays the high-resolution (1.63Å) crystal structure of human carbonic anhydrase (PDB 6pea), together with the structures predicted by RTF, ESM-2, and AF2. It is immediately apparent that all three algorithms predict this structure very well, displaying high pLDDT scores (Table 2). The RMSD values, relative to the crystal structure, are 0.76Å, 0.42Å, and 0.32Å for RTF, ESM-2, and AF2, respectively.

**Fig. 2.**
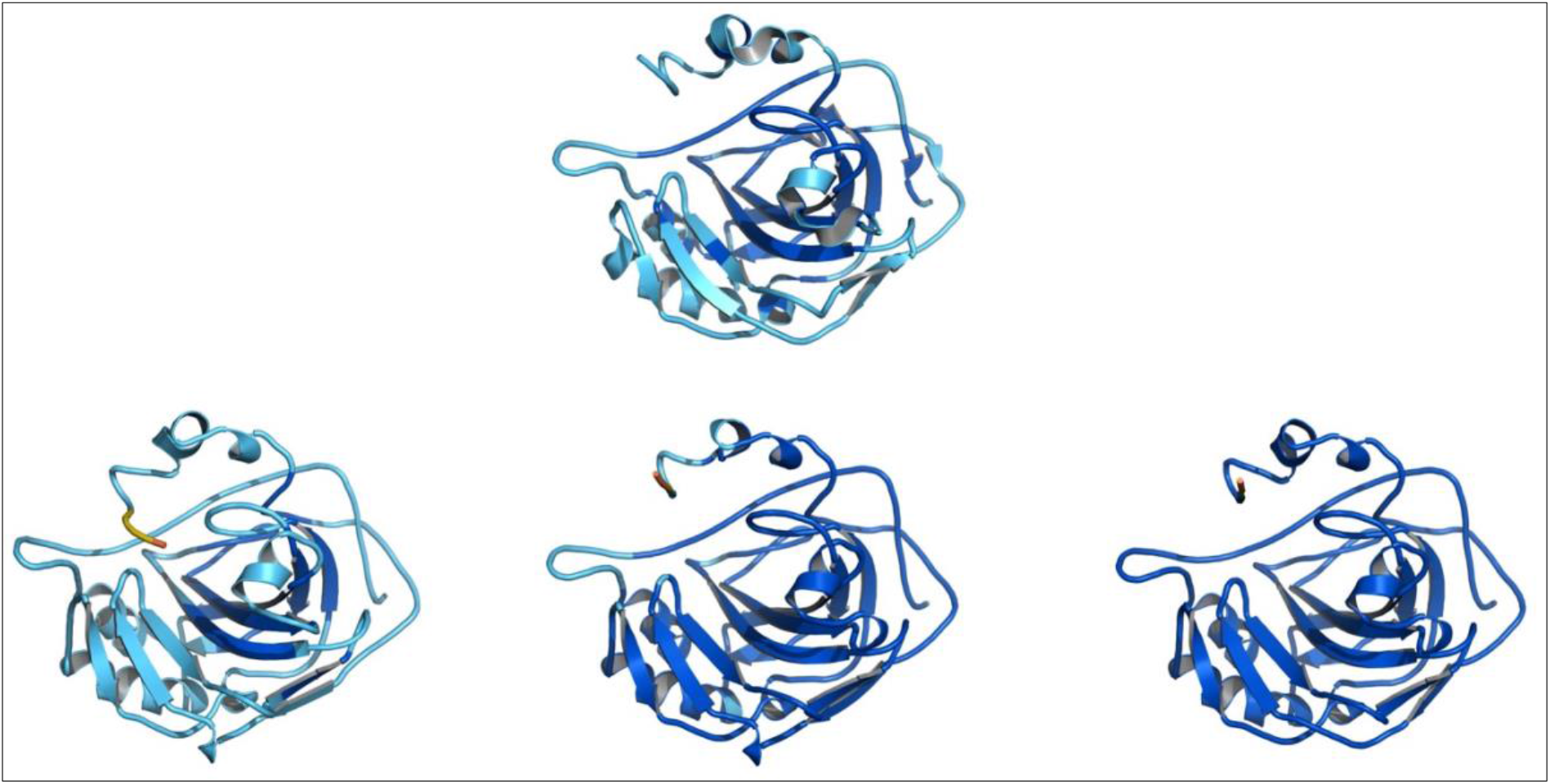
Crystal structure of human carbonic anhydrase, PDB 6pea (top), and structures predicted by RTF (left), ESM (center) and AF2 (right).

**Table 2.**
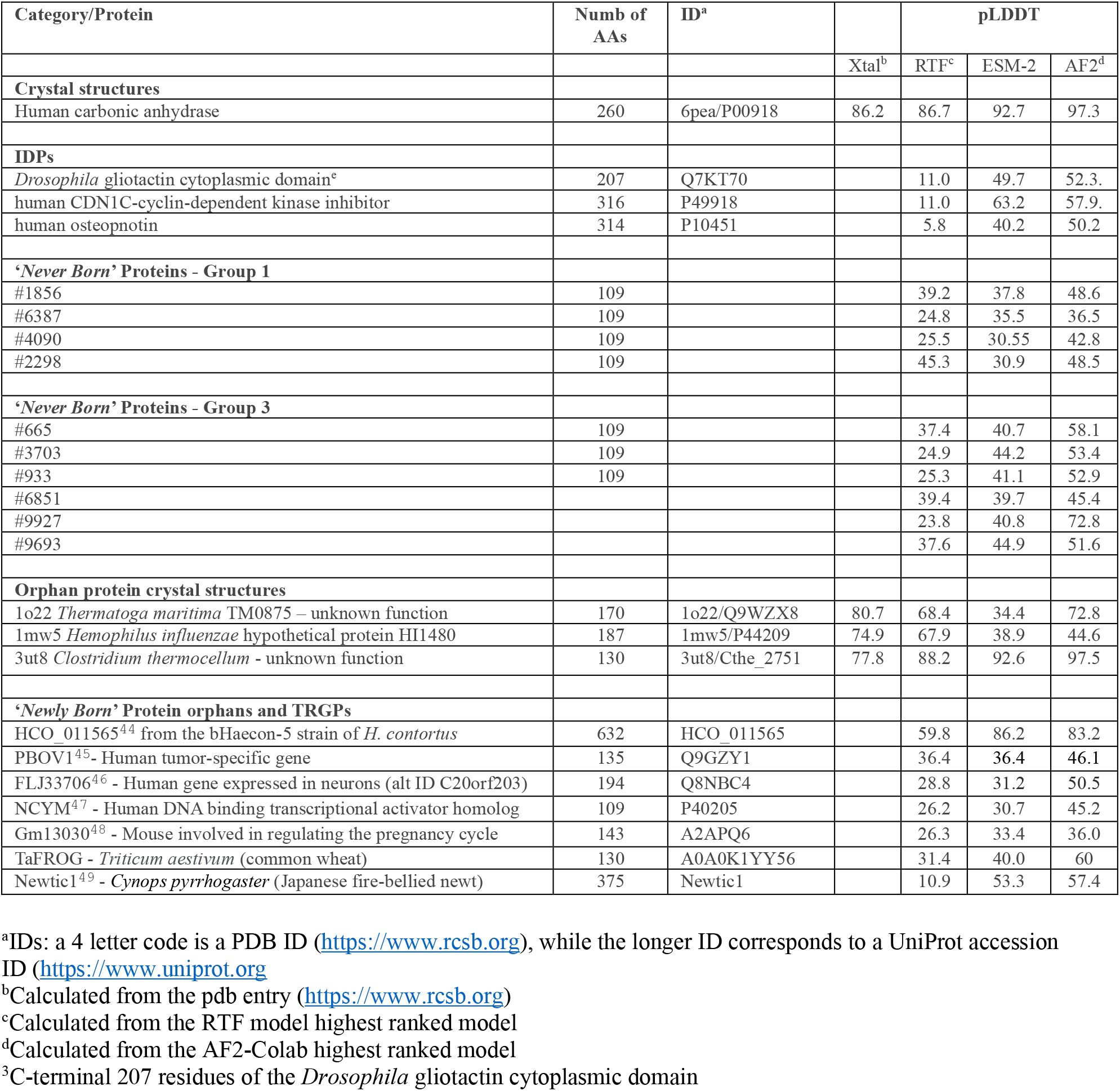

It is now well established that many native proteins are either partially or completely disordered and are known as intrinsically disordered proteins (IDPs)^36^. In a review by Uversky^58^ it is stated that “…eukaryotes typically have a higher disorder score than either archaea or prokaryotes, since 52-67% of eukaryotic proteins have long intrinsically disordered protein regions (IDPRs) (≥ 30 residues) as compared to 26-51% and 16-45% proteins with such long IDPRs in archaea and bacteria, respectively^59,60^.”

Several groups have examined how well AF2 can be used to predict IDPs^61-63^. We were also curious as to how well RTF and ESM-2 would predict their structures. Fig. 3 shows the predictions for three such proteins, the cytoplasmic domain of gliotactin, the ChE-like adhesion molecule (CLAM) from *Drosophila*^38^, human CDN1C-cyclin-dependent kinase inhibitor^39^, and human osteopontin^40^. RTF, ESM-2, and AF2 all three predict highly unfolded structures. However, in several of the models, substantial α-helical stretches are predicted, with their percentage in the RTF models being significantly higher than in those generated by the other two methods. These predicted helical sequences are different in the three models for each protein. In any case, they do not appear to correspond to real helical stretches since physicochemical data for the three proteins do not support their presence.

**Fig. 3.**
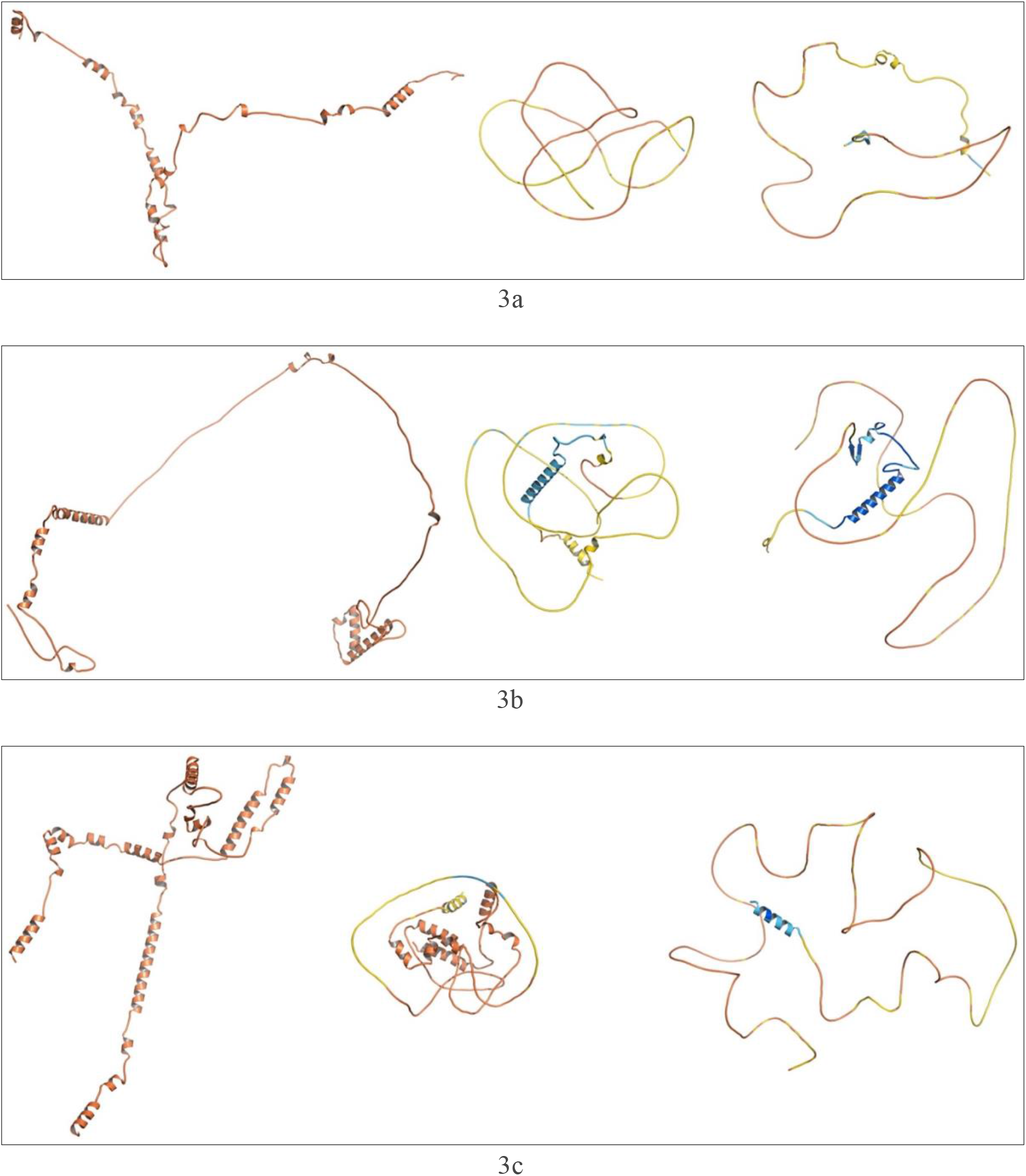
3D structure predictions for three IDPs using RTF, ESM-2, and AF2. A. Gli-Cyt, RTF (left), ESM (center),AF2 (right); b. CDN1C, RTF (left), ESM (center), AF2 (right); c. Osteopontin, RTF (left), ESM (center), AF2 (right).

As mentioned in the Introduction, Tretyachenko *et al*^8^ predicted, by use of bioinformatic tools, that some of the sequences that they subsequently expressed would be ordered/folded, with a high content of secondary structure elements, whereas others would be disordered/unfolded (IDPs), with a low content of secondary structure elements. They found that only low- significant matches were found by BLAST method for the whole set of random sequences^8^.

On the basis of their bioinformatic analysis, they selected groups of 15 proteins each from the random sequence library. Members of Group 1 were predicted to have high secondary structure, low disorder, and high solubility. Members of Group 3 were predicted to have low secondary structure, high disorder, and high solubility (Fig. 4). They showed that 31% of the proteins in group 1 expressed in soluble form, and used circular dichroism (CD) to demonstrate that they had substantial secondary structure. All the proteins in Group 3 expressed in soluble form, and had low secondary structure.

**Fig. 4.**
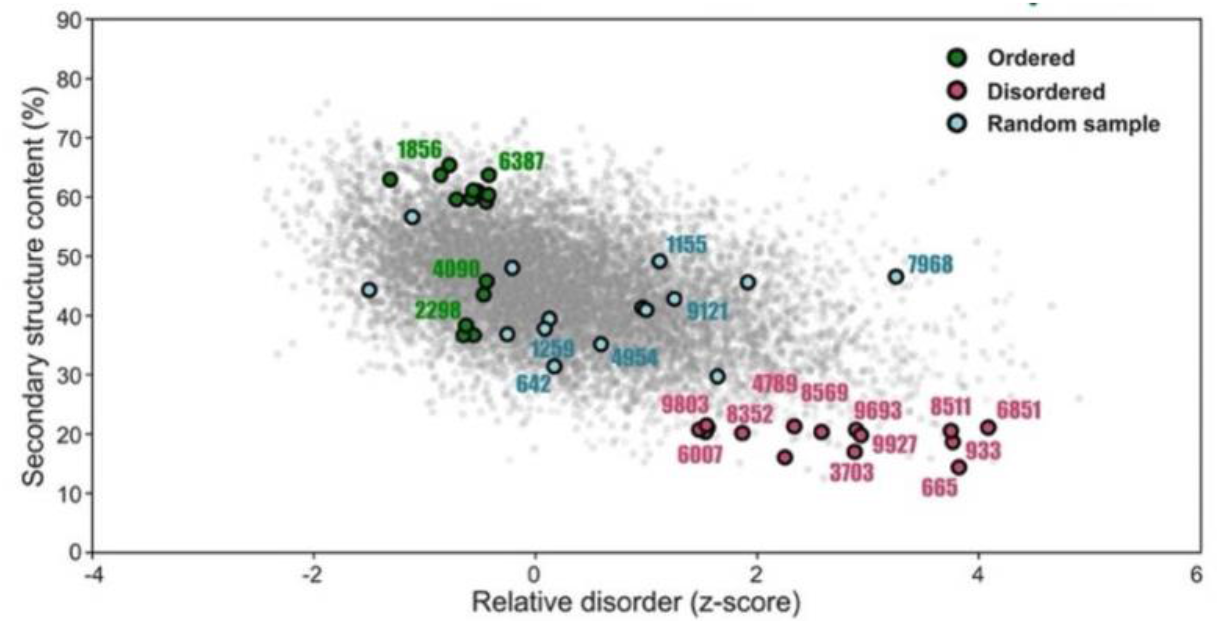
Selection of sequences from the set of ‘*Never Born’* proteins taken for experimental characterization. Secondary structure is plotted on the y-axis vs. relative disorder on the x-axis. Members of Group 1 (green circles) fall into the category of ordered/folded proteins, and members of Group 3 (red circles) fall into the category of disordered/unfolded proteins (IDPs) (taken with permission from the paper of Tretyachenko *et al*, 2017^8^).

We used all three structure prediction tools to model the structures of several members of Group 1 and Group 3 (Figs. 5 and 6). For all four members of Group 1, RTF predicts compact structures with a high percentage of secondary structure. The structures predicted by ESM-2 and AF2 are more open, with a higher percentage of disordered stretches. For Group 3, AF2 predicts very open structures, characteristic of IDPs; ESM-2 predicts somewhat more compact structures, and RTF predicts some of the structures to be open and others to be more compact.

**Fig. 5.**
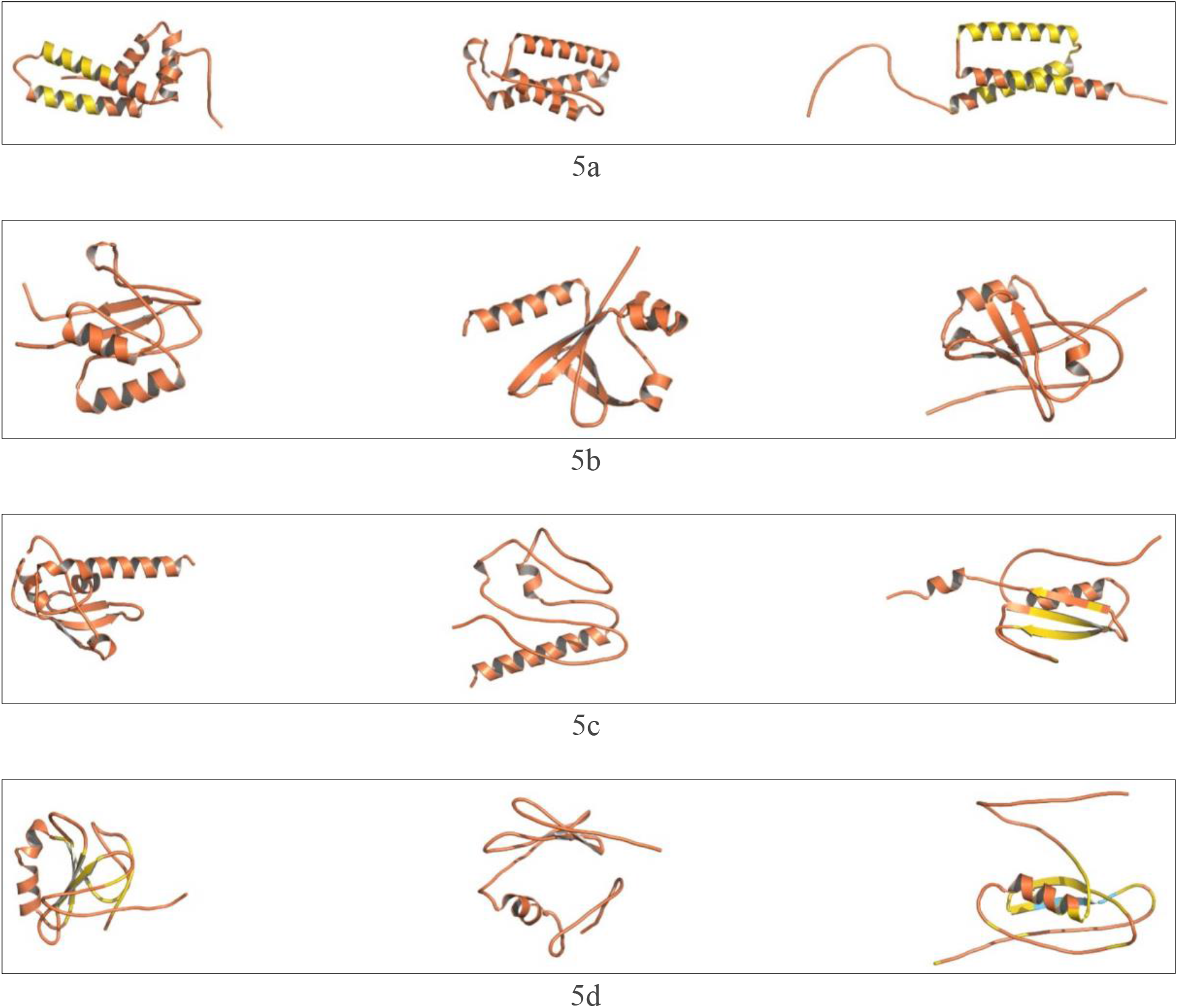
3D structure predictions for 4 members of Group 1 of *‘Never Born’* proteins, with RTF (left), ESM-2 (center), and AF2 (right) **a**. #1856; **b**. #6387; **c**. #4090; **d**. #2298.

**Fig. 6.**
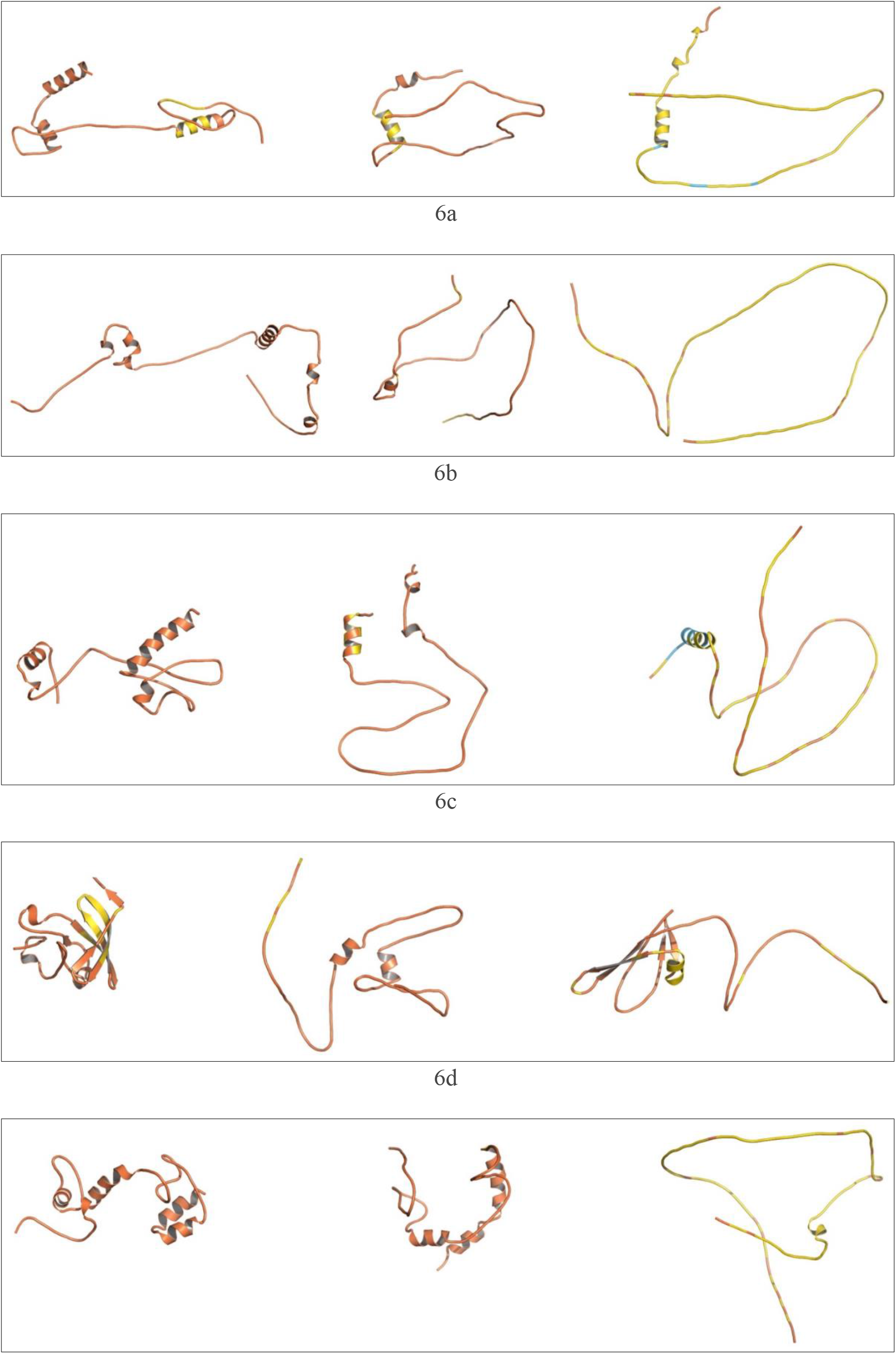
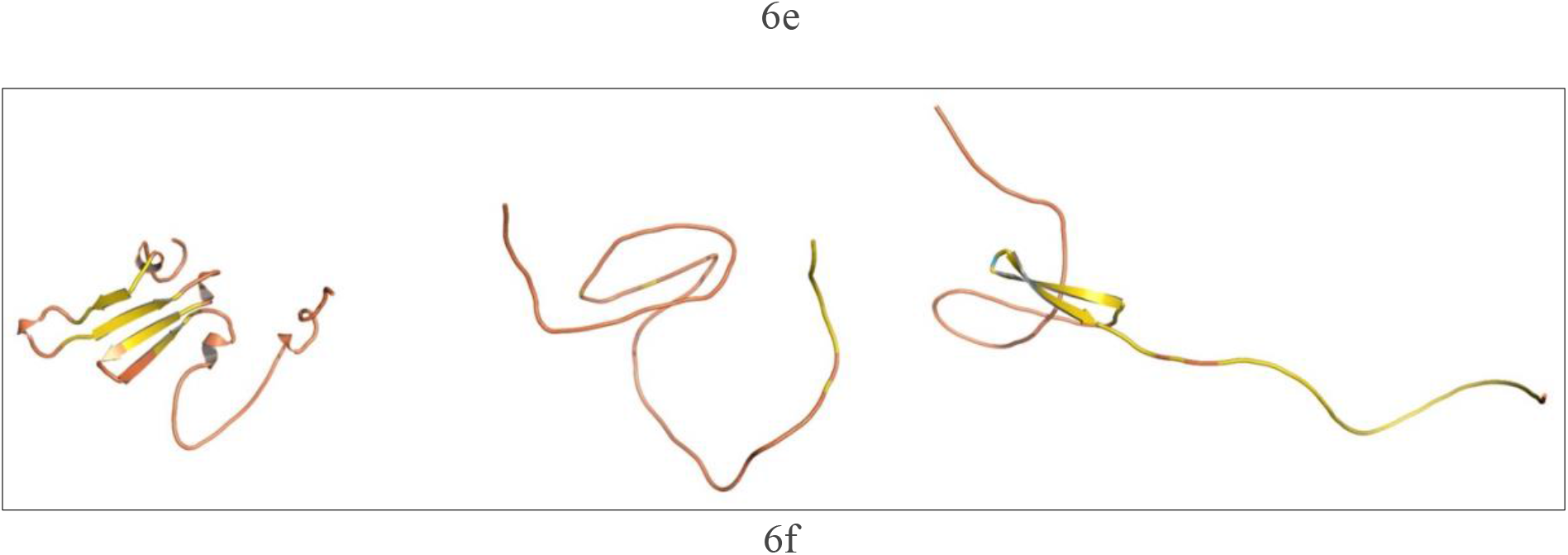
3D structure predictions for 6 members of Group 3 of *‘Never Born’* proteins., with RTF (left), ESM-2 (center), and AF2 (right) a. #665; **b**. #3703; **c**. #933; d. #6851; e #9927; f #9693.

Fig. 7 displays the accessible surface areas (ASAs) for the predicted models of proteins in Group 1 (red) and Group 3 (blue), using RTF, ESM-2, and AF2. A student T-test showed that all three predicted significant differences in ASA between Group 1 and Group 3, with p values being 0.046, 0.002, and 0.007 for RTF, ESM-2, and AF2, respectively.

**Fig. 7.**
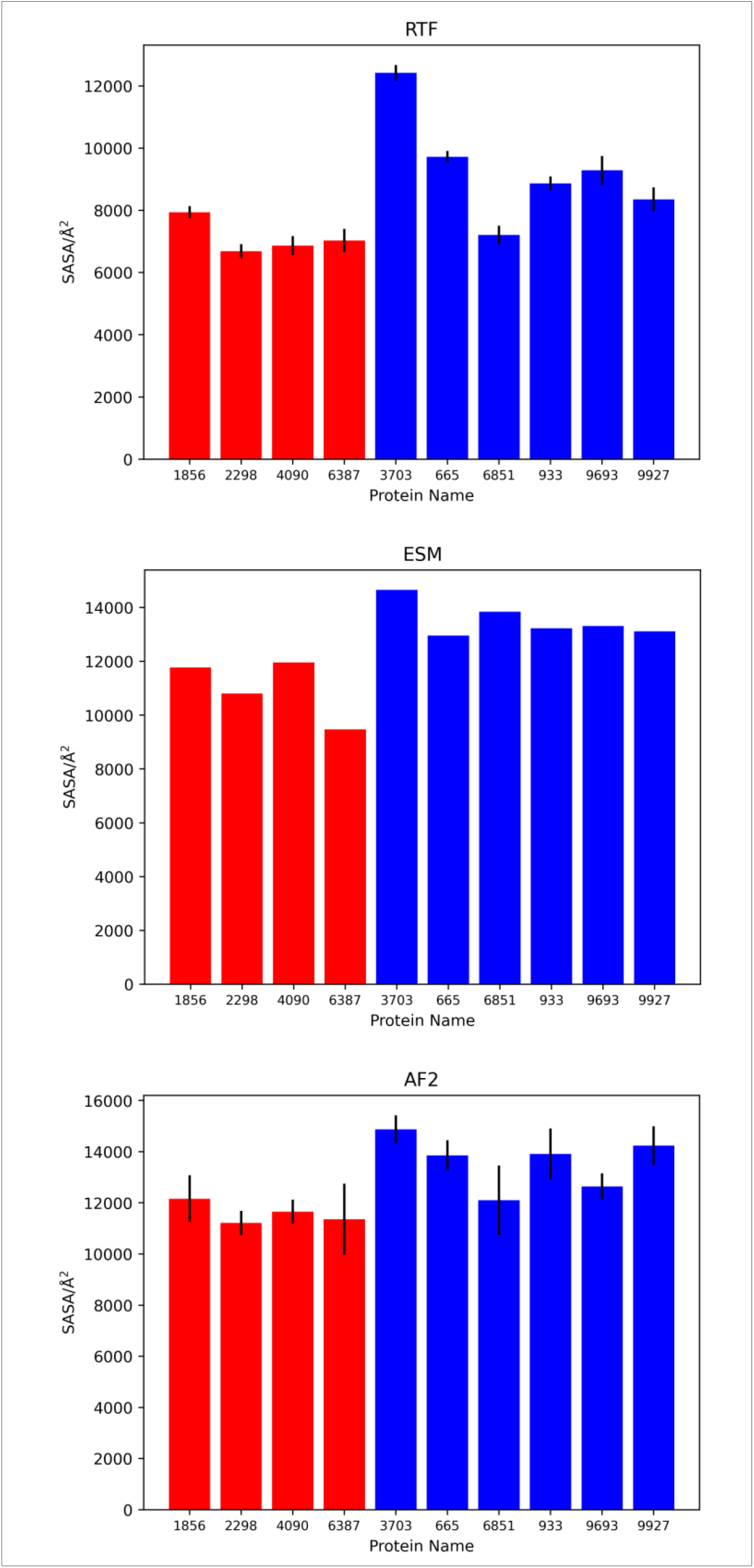
Comparison of the Accessible Surface Areas (ASAs) for Never Born proteins. ASA values for Group 1 are displayed in red, and those for Group 3 in blue. For RTF and AF2, the black vertical bars indicate the maximum and minimum values of the 5 predicted models. For ESM-2 no bars are shown, since it predicts only one model. Top panel, RTF; middle panel, ESM-2; bottom panel, AF2.

Despite the fact that research on orphan proteins is a hot topic, largely due to its evolutionary implications^64^, we were able to find only three crystal structures of orphan proteins in the PDB. Fig. 8a shows the crystal structure of orphan protein TM0875 from *Thermatoga maritima* (PDB 1o22)^41^, alongside predictions of its structure by RTF, ESM-2 and AF2. The RMSD values, relative to the crystal structure, are 1.37Å, 17.72Å and 1.62Å for RTF, ESM-2 and AF2, respectively. The crystal structure showed only 149 amino acids, although the sequence that was used for crystallization consisted of 170 amino acids. The pLDDT scores are well correlated with the RMSD values, with ESM-2 showing the lowest agreement with the X-ray structure. The authors pointed out that this was a novel fold, and application of the Dali ^54,65^ server reveals that it still maintains this status based on a much large number of experimental structures in the PDB. However, a BLAST search revealed many *Thermatoga* homologs, with the 15^th^ displaying 57% identity for 90% of the sequence. Thus, this protein is a TRGP rather than a true orphan.

**Fig. 8.**
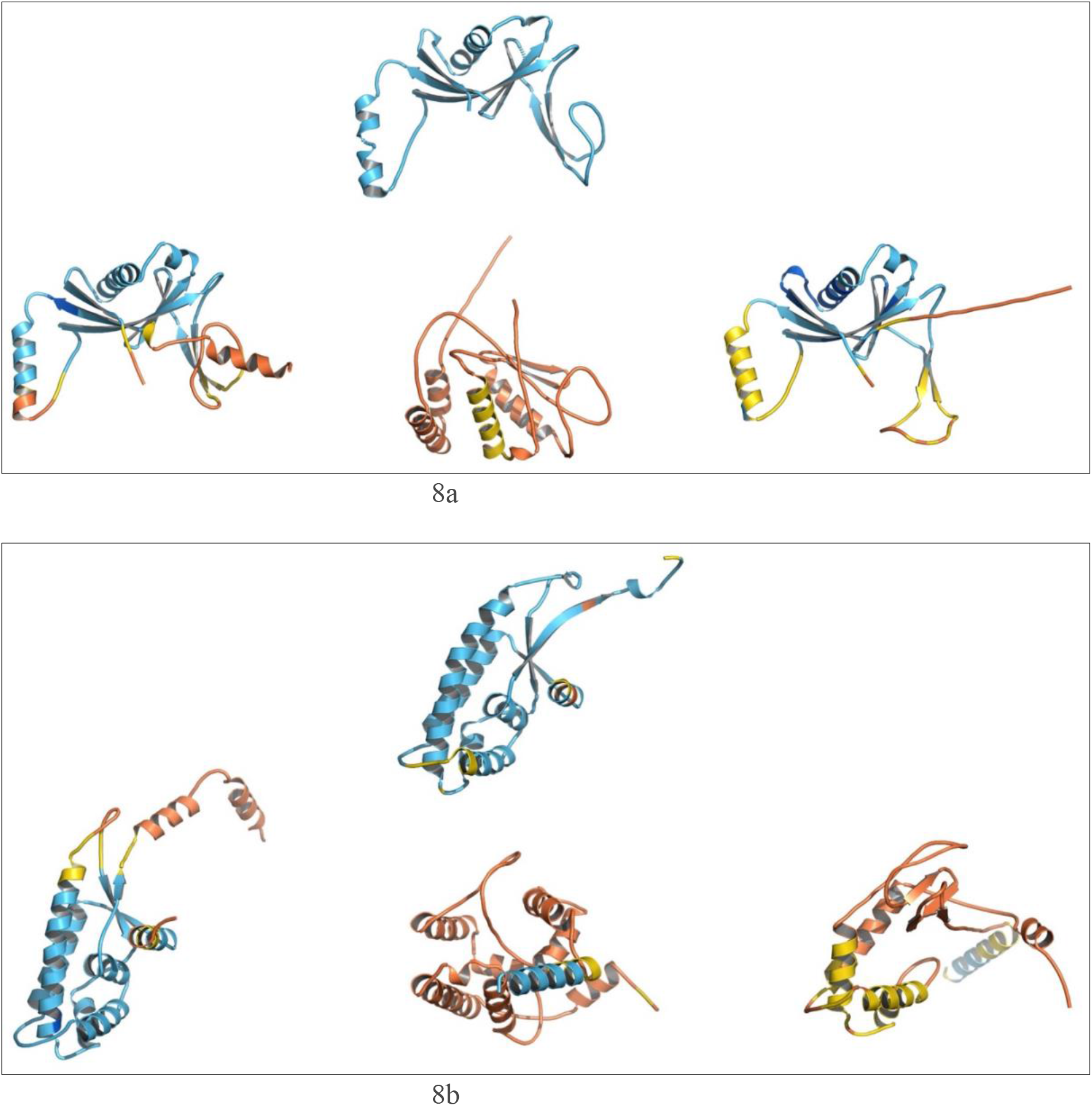
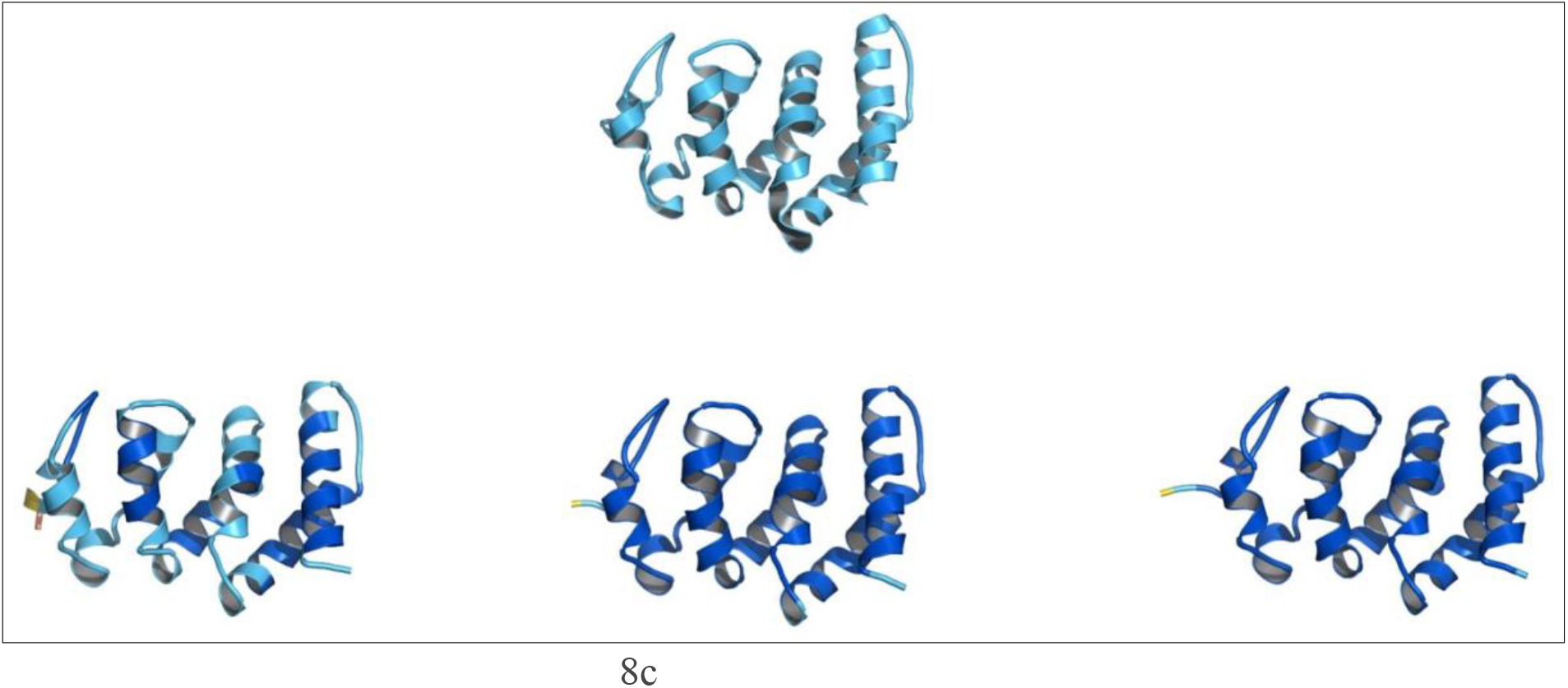
Cystal structures of an orphan proteins and two TRGPs, and the models generated by the three structure prediction algorithms. a) Crystal structure of the TRGP TM0875 from *Thermatoga maritima*, pdb 1o22 (top), and the structures predicted by RTF (left), ESM (center) and AF2 (right), b) Crystal structure of the orphan protein hypothetical protein HI1480 from *Haemophilus influenzae*, pdb 1mw5 (top), and structures predicted by RTF (left), ESM (center) and AF2 (right), c) Crystal structure of the TRGP, Cthe_2751, from *Clostridium thermocellum*, PDB- ID 3ut8 (top), and structures predicted by RTF (left), ESM (center) and AF2 (right).

Fig. 8b shows the crystal structure of a second orphan protein deposited in the PDB, that of the hypothetical protein HI1480 from *Haemophilus influenzae* (PDB 1mw5)^42^, alongside the structures predicted by RTF, ESM-2 and AF2. The RMSD values, relative to the crystal structure, are 1.31Å, 17.6Å and 7.3Å for RTF, ESM-2 and AF2, respectively. It is important to point out that the crystal structure was able to discern only 162 amino acids, although the sequence used for crystallization consisted of 187 amino acids. The pLDDT scores are well correlated with the RMSD values, with RTF being the only method showing good agreement with the X-ray structure, while AF2 showed some similarity. It is worth noting that, even though the 25-residue disordered region is not seen in the crystal structure, it is predicted to be largely helical by RTF. This is in keeping with our observation that in both authentic IDPs, and in members of Group 3 of the ‘Never Born’ proteins, helical stretches were quite frequently predicted by one or other of the three algorithms. In this case, too, the authors pointed out that this is a novel and unique fold, and application of the Dali server again reveals that it still maintains this status based on the much large number of experimental structures now available, as well as the entire AlphaFold Database. A BLAST search confirmed its orphan status.

Cthe_2751, whose crystal structure has been determined [PDB-3ut8], is a protein from *Clostridium thermocellum* with unknown function, which had been reported to be a singleton^43^. However, a BLAST search that we performed revealed many homologs from the genus *Clostridium*. So, in fact, it is not an orphan, but rather a TRGP. Examination of its crystal structure revealed an all α-helical topology similar to those observed for nucleic acid processing proteins^43^ (Fig. 8c).

We thought that it would be of interest to use RTF, ESM-2 and AF2 to predict the structures of orphan proteins and TRGPs for which no experimental structures were available, both in order to see whether they would predict novel folds, and to find out how the predictions of the three algorithms would compare, To this end we selected seven proteins whose characterization as orphan proteins, or as TRGPs, appears to be well established, which have been shown experimentally to have defined biological functions, and for which the necessary sequence data are available. These proteins are:

- HCO_011565 from the Haecon-5 strain of the nematode, *Haemonchus contortus*. This is a 632-residue protein that has been expressed and characterized by Taki *et al*.^44^, who showed that it was the target of a nematodicidal small molecule. They also modeled it using AF2, which predicted that it has a well-defined 3D structure, with a transmembrane domain at its C-terminus. When we performed BLAST on this protein, we found that the first 74 homologs retrieved were all from nematodes. The 75^th^ was from a tick, with the E value increasing from 1xe^-9^ to 3xe^-6^ between the 74^th^ and 75^th^ homologs. Thus, it is clearly a product of a TRG rather than a true orphan.
- PBOV1 is a human *de novo* gene with tumor-specific expression that is associated with a positive clinical outcome of cancer^45^. It is highly expressed in primary gliomas and breast tumors. PBOV1 codes for a protein that contains 135 residues, whose function is unknown. The authors reported that they were unable to find any orthologs, whether in humans or in any other species. Our BLAST search produced many hits, but all were described as “Low Quality Proteins”, meaning that it is uncertain if they are, in fact, real proteins So, the PBOV1 gene product appears to be a true human orphan protein.
- FLJ33706 (alternative gene ID C20orf203) was identified as a *de novo* human gene in studies concerned with nicotine addiction^46^. It is abundantly expressed within neurons in several areas of the human brain, including cortex, cerebellum and mid-brain, and elevated expression was observed in Alzheimer’s disease brain samples. It codes for a protein that contains 194 residues, and the authors identified it as human-specific. Our own BLAST search also revealed no orthologs. So, it, too, expresses a true human orphan protein. As for PBOV1, our BLAST search for FLJ33706 produced many hits, but all were described as “Low Quality Proteins”, again meaning that it is uncertain if they are, in fact, real proteins So, the FLJ33706 gene product, too, appears to be a true human orphan protein.
- NCYM, a cis-antisense gene of the MYCN oncogene, encodes a *de novo* evolved protein that regulates the pathogenesis of human cancers, especially neuroblastoma*s*^47^. The NCYM protein, which contains 109 residues, inhibits the gsk3β kinase, which, in turn, promotes degradation of the MYCN gene product. So NYCM is the first *de novo* evolved protein to act as an oncopromoter in human cancer. For NCYM, our BLAST search revealed many “Low Quality Homologs” and a few genuine hits, all from monkeys. So, the gene product appears to be TRGP.
- Gm13030, a protein encoded by a young protein-coding gene, containing 143 residues, is specifically expressed in the oviduct of the female mouse^48^. If the gene expressing it is knocked out, the pregnancy cycle is shortened, and the infanticide rate increased^48^. In this case, too, our BLAST search revealed no orthologs. So, the gene product appears to be a true protein.
- TaFROG, which stands for *Triticum aestivum Fusarium* resistance orphan gene, expresses an orphan protein that confers resistance on wheat (*T. aestivum*) to the mycotoxigenic fungus, *Fusarium graminearum*^*49*^. The TaFROG protein, which contains 130 residues, and was found to be an IDP, is localized to the nucleus, and acts by interacting with the α subunit of the sucrose non-fermenting-related kinase 1. Our BLAST search revealed two uncharacterized proteins from related wheat, and a few hypothetical proteins. Thus, the TaFROG protein appears to a TRGP with a small number of identified homologs.
- Newtic1 is an orphan protein expressed in the regenerating limbs of the adult Japanese fire-bellied newt, *Cynops pyrrhogaster*^*50*^. It is found in only one other closely related newt. It is specifically expressed in a subset of erythrocytes that form clumps that accumulate in the distal portion of the regenerating limb, and may be involved in clump formation. It contains 375 amino acids, with a transmembrane sequence near its N-terminus. Our BLAST search did not reveal any additional homologs than the one referred to already. So, the gene product appears to be a true orphan protein.

The number of amino acids for these seven orphan/TMG proteins ranges from 109 for NCYM to 632 for HCO_011565, and their pI values range from 4.99 for Newtic1, which contains 7.7% Glu and 5.1% Asp residues, with a net charge at pH7.4 of -15.67, to 12.23 for FLJ33706, which contains 10.3% Arg and 2.6% Lys residues, with a net charge at pH7.4 of +15.25

As an initial step in characterizing these seven proteins, we utilized FoldIndex^56^ and flDPnn^57^ to investigate whether they were predicted to be intrinsically disordered or folded. Although these algorithms make similar predictions, since flDPnn was selected as the best disorder predictor in the first Critical Assessment of Protein Intrinsic Disorder Prediction (CAID)^66^, the data displayed in Fig. 9 were generated using it. Four of the proteins are predicted to be almost completely folded, although FLJ33706 has a short, disordered stretch, in the middle of its sequence. The other two, TaFROG and Newtic1, are classified as IDPs, since they are predicted to be disordered throughout almost their entire sequences.

**Fig. 9.**
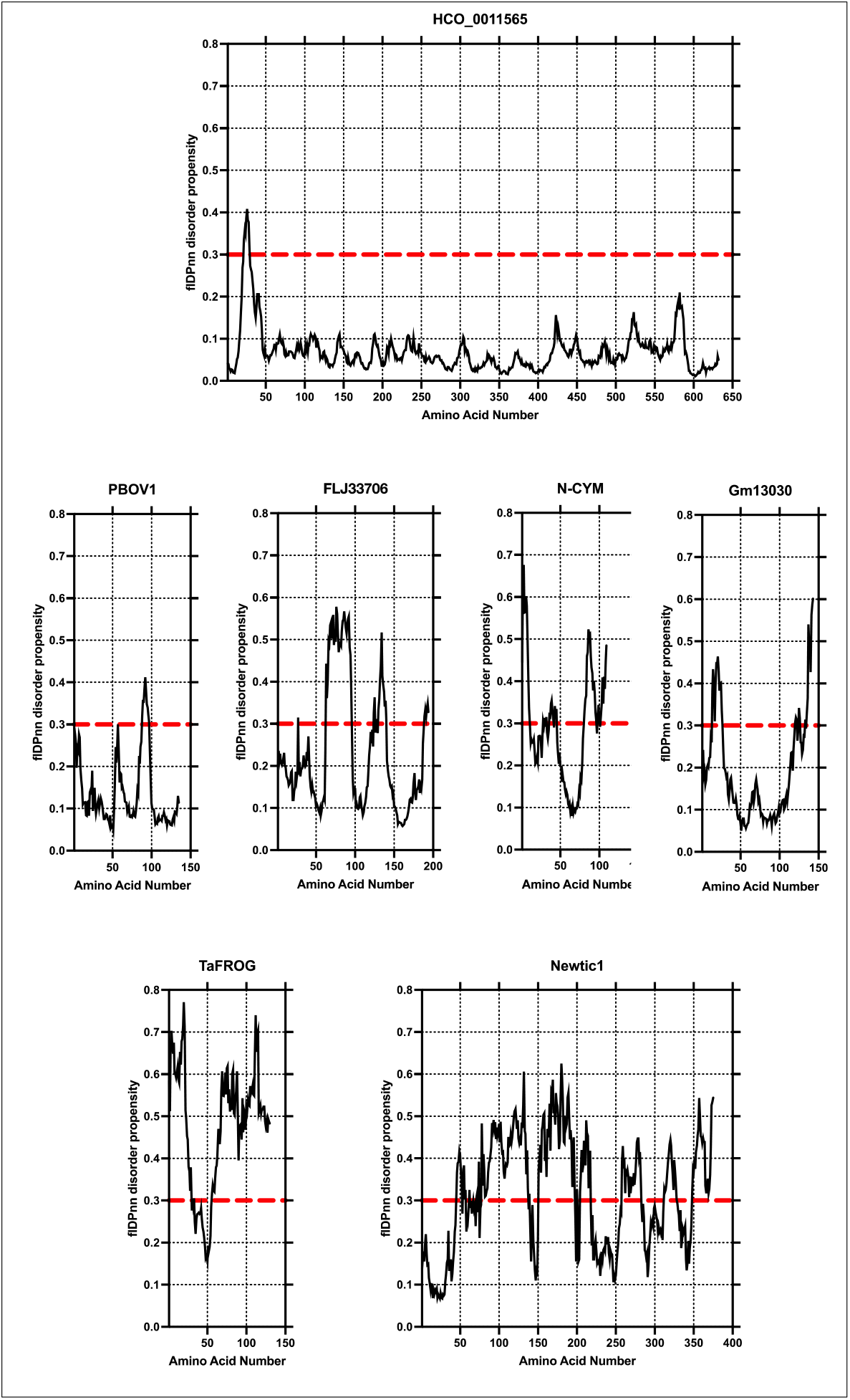
flDPnn predictions of order/disorder in seven well-studied orphan or TRG proteins. (a) HCO_011565; (b) PBOV1; (c) FLJ33706; (d) NCYM; (e) GM13030; (f) *Ta*FROG; (g) Newtic1. Sequences above the red horizontal line are classified as disordered, and below the line as ordered.

Of the seven structures generated using the three structure prediction algorithms, only the first, that of HCO_011565, shows fully folded and almost identical structures (Table 3), with high pLDDT scores (Table 2). Most likely, this is for two reasons. Firstly, rather than being a *true* orphan, HCO_011565 is the product of a TRG^44^, with the BLAST search having revealed that the first 74 homologous sequences with the lowest E values, were all from nematodes. Secondly, the DALI server revealed a number of hits for the entire predicted structure, as well as for the three sub-domains predicted by all three algorithms.

**Table 3.** Comparison of the RMSD values for the 3D structures of HCO_011565 predicted by the three algorithms.

| Prediction Algorithms | RMSD (Å) | Number of atoms used |
| --- | --- | --- |
| ESM-2 vs RTF | 3.5 | 3,640 |
| AF2 vs RTF | 3.6 | 3,782 |
| AF2 vs EMS-2 | 1.2 | 3.773 |

The alignments were performed with PyMOL super command, which automatically determines the number of atoms to be used.

The flDPnn disorder algorithm predicted that TaFROG^49^ and Newtic1^50^ are mostly unfolded, and indeed all three structure prediction algorithms produce very open structures for both of them. It is interesting that in Newtic1 a substantial stretch near its N-terminus is predicted to be ordered. Casco-Robles *et al*.^50^ predicted that the sequence F^17^LWALMSTASMVSTLVALLLLCGLC^40^, is a transmembrane sequence; it is therefore likely to be α-helical, and such an assignment is, indeed, made by all three structure prediction algorithms (Fig. 10).

**Fig. 10.**
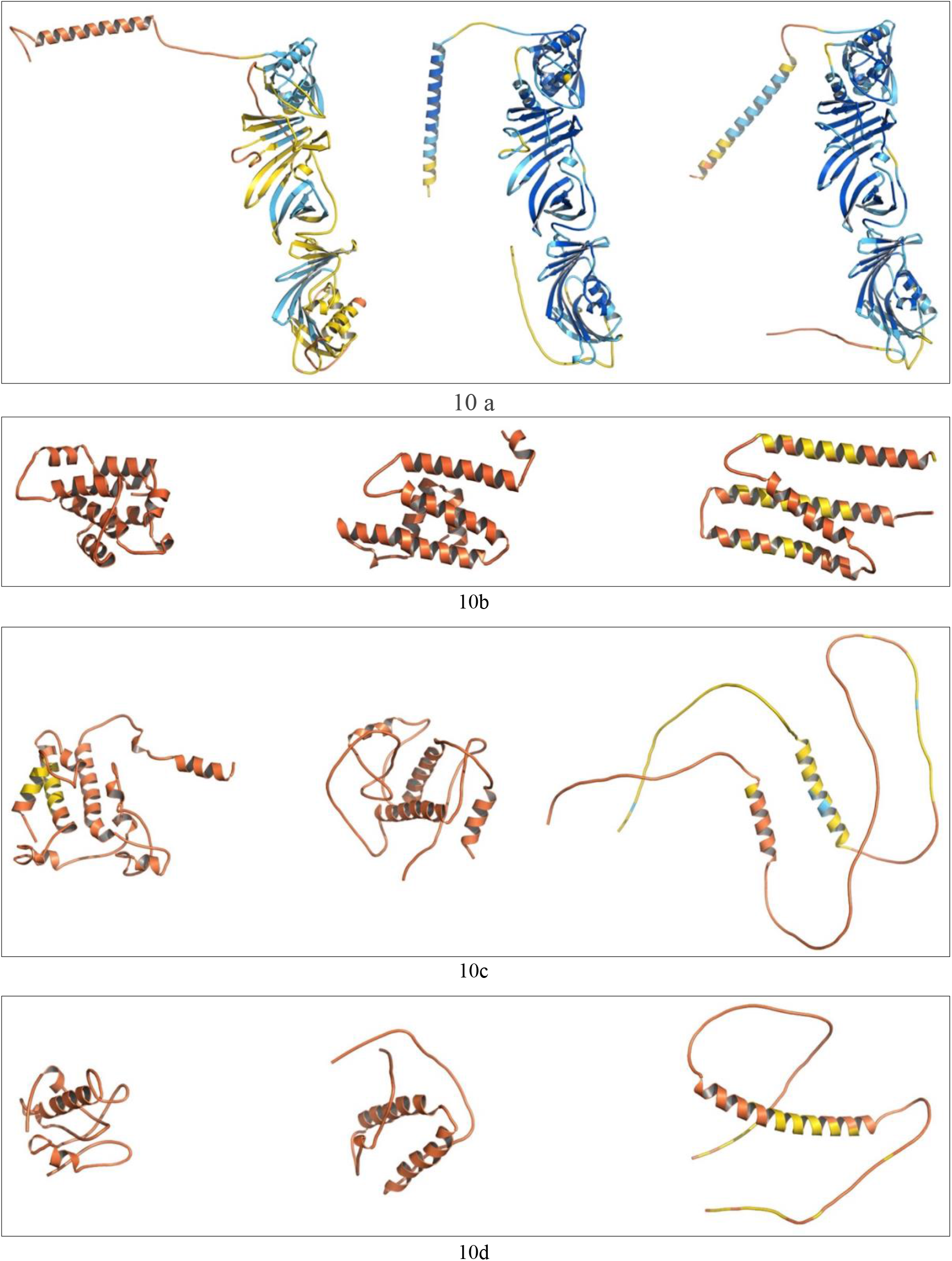
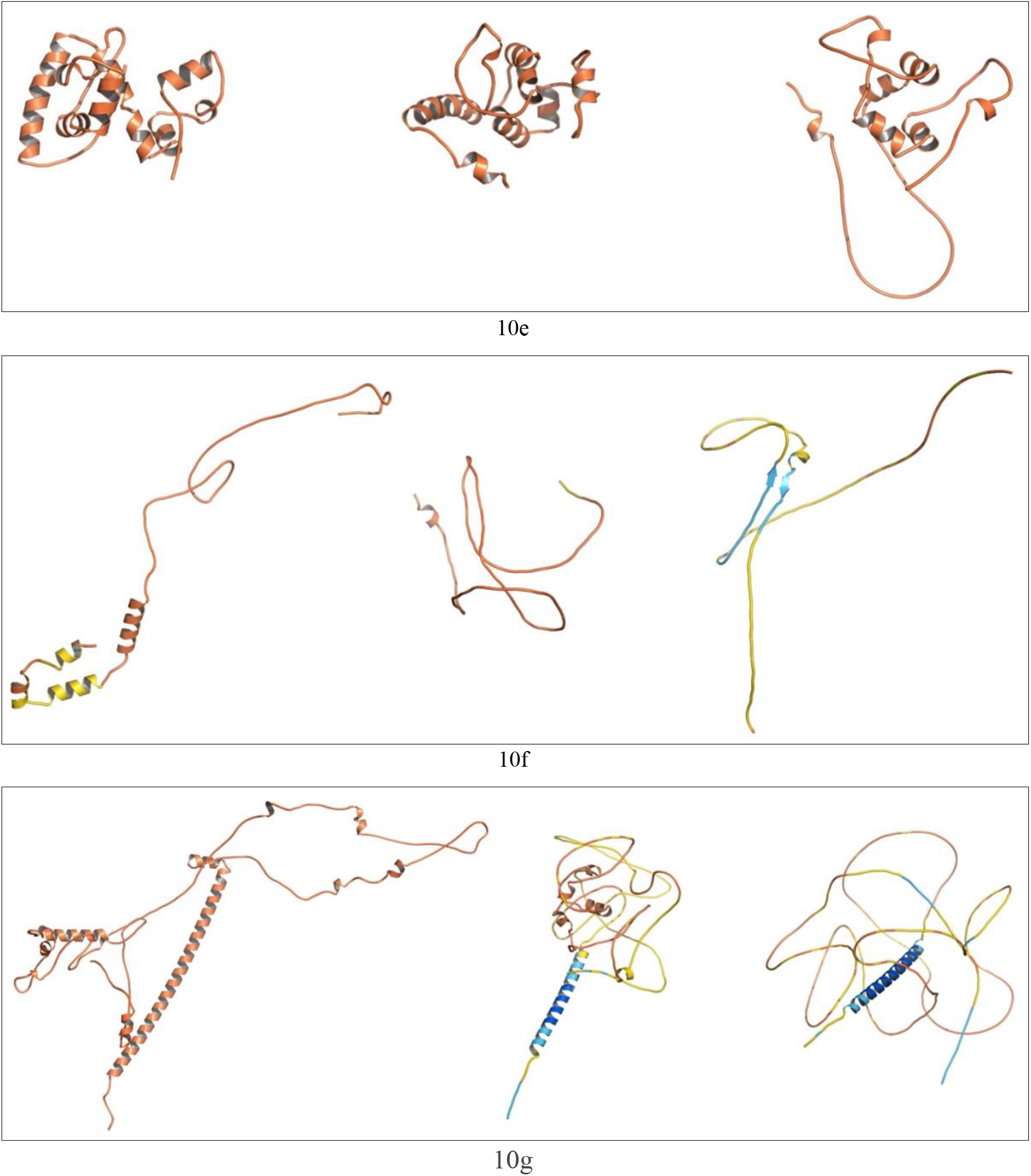
3D structure predictions for 7 orphan/TMG proteins using RTF (left), ESM (center), and AF2 (right). **a**. HCO_011565; b. PBOV1; **c**. FLJ33706; d. NCYM; **e**. Gm13030; f. *Ta*FROG; g. Newtic1.

The remaining four proteins^45-48^ are predicted to be predominantly folded by flDPnn. However, the three structure prediction algorithms yield quite different results for each of them. We should, of course, stress that the pLDDT scores are very low for all these structures, using all three algorithms. However, our present interest is not in their precise 3D structures, but rather in their overall folds. We think, therefore, that the information that we present, and our interpretation of it, is meaningful. For all four proteins, RTF yields the most compact structures, with a high percentage of secondary structure (mostly α-helical), while for the structures predicted by ESM-2, there is substantial secondary structure, but the structures are somewhat less compact. In the case of AF2, three of the structures are very open, but that of PBOV1 is quite compact, and shows some similarity to that predicted by the other two algorithms. As discussed above, for the three orphan proteins or TRGPs for which the crystal structures have been determined to high resolution, RTF predicts all three very well, AF2 predicts two quite well (PDB-ID 1o22 and PDB-3ut8) and the other relatively poorly (PDB-ID 1mw5), while ESM-2 only predicts one very well (PDB-ID 1ut8).

To return to consideration of HCO_011565, apparently, as consequence of its having many close homologs, both RTF and AF2 make predictions with very high pLDDT scores. Furthermore, ESM-2, which does not use multiple sequence alignment (MSA), also predicts a structure with a high pLDDT score. Visual inspection shows that all three predicted structures are very similar (Fig. 10a), and indeed, the RMSD values are very low (Table 3). It is interesting that AF2 and ESM-2 give essentially identical structures, although the two algorithms are quite different.

## 4. DISCUSSION

**The discovery of ‘Newly Born’ orphan proteins**^**23-26**^, **and of proteins coded for by TRGs, viz**., **TRGPs, which utilize DNA sequences that were previously not expressed, raises cogent questions as to how the polypeptides coded for by such sequences evolve into biologically active proteins**.

The principal goal of this study was to find out if the powerful structure prediction algorithms that have emerged in the past three years would be useful for structure analysis of such emergent proteins. AF2 and RTF use MSA of homologous proteins as an important element for structure prediction^12,31^, while ESM-2 does not^32^.

True orphan proteins have no sequence homology to any existing protein. We thought, therefore, that the ‘*Never Born*’ proteins generated and investigated by Tretyachenko *et al*^8^ would serve as a valuable benchmark for comparison. These authors expressed and purified a substantial number of polypeptides that had compositions similar to those of authentic proteins but whose sequences were random. Based on CD measurements, they found that a significant number of the random sequences, Group 1, were compact, with substantial secondary structure, while another set, Group 3, appeared to be IDPs. Indeed, the sequences of the Group 3 members were very similar to those of well characterized authentic IDPs. We examined members of these two groups of polypeptides using all three structure prediction algorithms. Although the structures predicted differed significantly in detail, and had low pLDDT scores, by analysis of the ASA data, it was clear that the members of Group 1 were significantly more compact than those of Group 3 (Fig. 7). Moreover, for members of Group 1, both RTF and ESM-2 predict significantly more compact structures than AF2. For Group 3, for which both physicochemical evidence and IDP algorithms indicated that they were intrinsically disordered^8^, all three structure prediction algorithms predicted remarkably well that they were disordered, with one exception, #6851 in the case of RTF (Fig. 6d). It would be interesting to determine the structures of members of Group 1, to see if their folds correspond to one or the other of those predicted by the three algorithms, or to some other folds, whether novel or preexisting.

We similarly analyzed ‘Newly Born’ orphan proteins and TRGPs. Parenthetically, some of the proteins we examined, which had been defined as orphans in the cited publications, turned out to be TRGPs.

As presented under Results, screening of the PDB revealed only three ‘orphan’ proteins for which crystal structures had been deposited. Only one of these was a true orphan, PDB-ID 1mw5. The other two, PDB-ID 1o22 and PDB-ID 3ut8, were TRGPs. The first two had novel folds, while the third, 3ut8, had a fold that had already been described in proteins that were functionally completely unrelated. It should be noted that the models predicted for this structure by all three algorithms are in excellent agreement with the crystal structure, very much better than the predictions made for the other two orphans/TMGPs displayed and discussed above. This may be attributed to the large number of homologs revealed by BLAST and possibly due to the fact that structures of proteins with similar folds are present in the PDB. Thus, perhaps not surprisingly, not all orphan proteins, or TMGPs, display novel folds.

For the seven orphans, or TRGPs, for which no 3D structure data were available, the disorder predictor, flDPnn, indicated that 5 were compact and 2 were IDPs.

For one of the compact structures, HCO_011565, the three structure algorithms predicted remarkably similar structures with very high pLDDT scores. In contrast, for the other four, all three algorithms predicted structures with very low pLDDT scores. It is plausible that the high quality and similarity of the structures predicted for HCO_011565 is due to the fact that the DALI server identified many structures in the PDB with similar folds for the entire protein and for each of its three domains. In addition, BLAST identified many homologous nematode sequences. The quality of the data is similar to that of the data obtained for 3ut8, for which, in addition to the existence of homologous sequences detected by BLAST, and of proteins from other families that displayed the same fold that DALI detected, the crystal structure was available to confirm the predictions.

Inspection of the predicted folds for both Never Born and Newly Born proteins shows several in which 3 or 4 α-helices appear to be aligned, notably the Never Born #1856, and the Newly Born PBOV1. The Dali server revealed good matches with 3- and 4-helix bundles in several proteins in the PDB.

Monzon *et al*.^67^ recently examined 250 protein sequence families in the AntiFam resource^68^, which are thought to be *spurious* proteins. Specifically, they are believed to be ORFs either on the opposite strand or in a different, overlapping reading frame, with respect to the true protein-coding or non-coding RNA gene^68^. Monzon *et al*.^67^ conjectured that proteins belonging to these families would not fold into well-folded globular structures. Using AF2, they confirmed this prediction with one exception. To the best of our knowledge, these spurious protein sequences were not examined with RTF or ESM-2. Since some of the data presented above suggest that AF2 may over-predict sequences lacking homologs to be unstructured, it would be worth using these two alternative algorithms to see whether they would predict that some of these sequences might indeed fold into compact structures. Thus, they may not be spurious and possibly have biological functions that could be tested experimentally.

In the Introduction, we referred to a recent study from the Baker lab^13^, in which 129 random sequences were modeled with RTF. These models, presumably with low reliability scores, served as useful starting models for optimization into folded proteins. Thus, although these models have low pLDDT scores, and thus most likely differ significantly in detail from the actual structures, they can provide good low-resolution starting points to evolve in the lab into well-folded and biologically active structures.

One conclusion that can be drawn from the data presented above is that some orphan proteins and TRGPs display novel folds that do not overlap with folds already present in the PDB. Indeed, the total number of distinct protein folds has been the topic of heated controversy^69,70^. In the present study, novel folds are observed for two of the three orphan proteins for which crystal structures exist, *i*.*e*., 1o22 and 1mw5 (Figs 8a and 8b).

One of the major reservations that has been made with respect to orphan proteins being ‘*Newly Born*’ proteins, coded for by sequences that were previously non-coding sequences, is that the genes in question might have undergone such rapid evolution that their homology to their predecessors was no longer recognizable^71,72^. The fact that two of the orphan proteins studied here display novel folds substantially weakens this argument.

In retrospect, it is not surprising that many random polypeptide sequences of a suitable amino acid composition, at a first approximation with a high content of hydrophobic residues and a low net charge^73^, will yield a compact structure containing substantial secondary structure motifs, as shown by Tretyachenko *et al*.^8^. This may be due to the fact that the sequences retained the amino acid compositions of the natural proteins from which they were generated, thus not being completely random.

The paradigm change introduced by Kuwajima and by Ptitsyn in the 1980s^74,75^ resulted in the realization that the newly synthesized polypeptide that emerges from the ribosome does not persist as an extended unfolded polypeptide unless it is an IDP, but rather collapses to what is termed a ‘Molten Globule’ (MG), a compact structure somewhat larger than the fully folded native structure (Fig. 11). The MG contains substantial secondary structure elements, but lacks the precise tertiary interactions of the native structure. Small proteins may spontaneously undergo a transition to the native state, whereas larger proteins may require the assistance of molecular chaperones to complete the folding process. The spectroscopic data of Tretyachenko *et al*.^8^ only tell us that compact structures, with secondary structure elements, have been produced by their random polypeptide sequences. When a native protein unfolds to a MG or some other partially unfolded species, hydrophobic amino acid side chains buried in the hydrophobic core become exposed. The degree of their exposure can be checked by use of the amphiphilic probe, 1-anilinonaphthalene-8-sulfonate (ANS), whose fluorescence is enhanced upon interaction with the hydrophobic residues^76^. It would, therefore, be interesting to compare the ANS fluorescence of the *‘Never Born’* proteins generated by Tretyachenko *et al*.^8^ to that of typical globular proteins in their native state. It is worth mentioning that in an early study on the folding of polypeptides with random sequences of simplified amino acid composition, NMR data indicated loose packing of the folded state^9^.

**Fig. 11.**
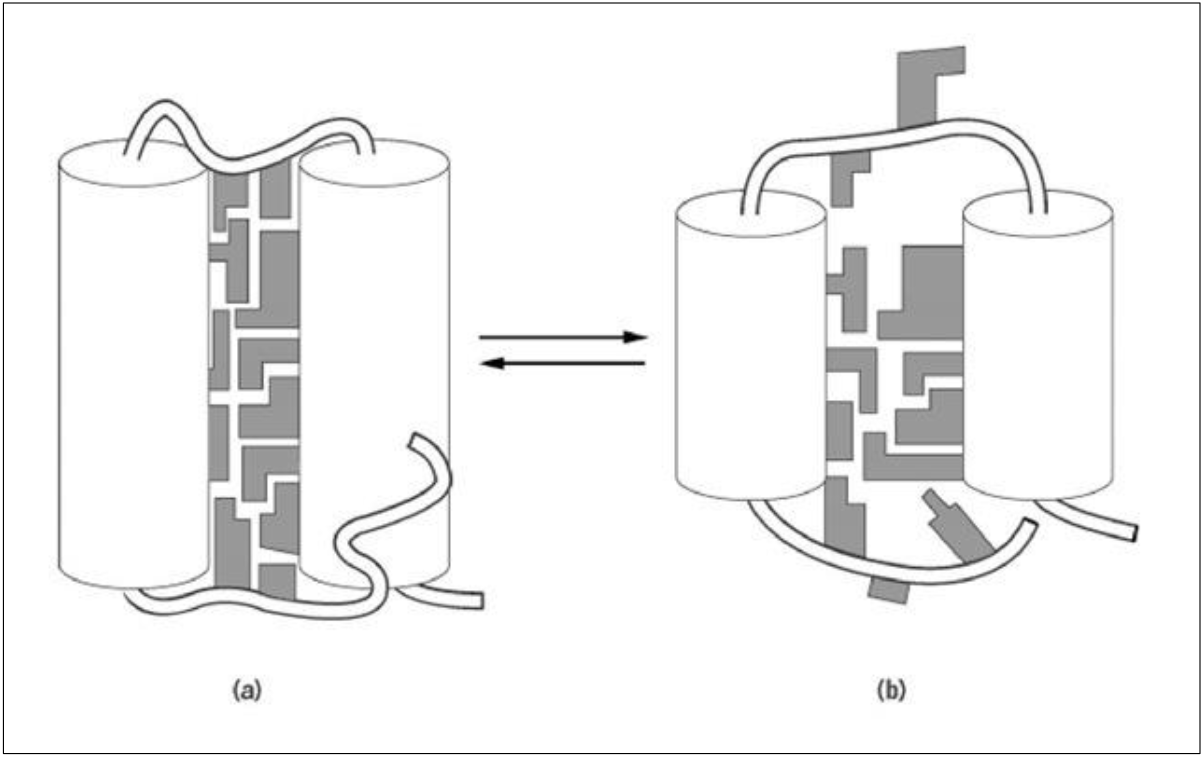
A schematic model of the native (a) and molten globule (MG) (b) states of a protein molecule. For the sake of simplicity, only two α-helices are represented. According to this model, the MG preserves the mean overall structural features of the native protein, but differs from the native state mainly in being more loosely packed, and thus having a volume larger by *ca*. 10%, and exposing more hydrophobic surfaces (reproduced from http://what-when-how.com/molecular-biology/molten-globule-molecular-biology). This figure was previously published in Tretyachenko *et al*., Sci Rep. 2017;7(1):15449 licensed under a Creative Commons Attribution 4.0 International License http://creativecommons.org/licenses/by/4.0/ and reproduced here with no change.

In any event, one can speculate that ‘*Newly Born*’ proteins might, initially, assume a MG-like conformation that would resemble that of the ‘*Never Born*’ proteins, and that mutations, coupled with natural selection, might convert some of them into ‘native’ orphan proteins with novel biological activities. This may be considered analogous to what occurred in the study in which the hallucinatory proteins were generated^13^.

Why do RTF and ESM-2 do relatively well in predicting plausible compact structures for randomized sequences that have been shown to be compact experimentally, whereas AF2 often makes predictions that are either clearly wrong or implausible? In a recent brief survey of the principles underlying AF2^77^ it was emphasized that it makes extensive use of the detection of conserved interactions of residues that are remote from each other in the linear sequence. This approach was earlier proposed, and implemented with a certain degree of success by Marks and Sander^78^. Obviously, such conserved interactions of distant residues would not exist in randomized sequences. Nor would such conserved interactions be available in orphan proteins, which lack ancestral homologs and, furthermore, in some cases display novel folds. Apparently, even if RTF makes use of such conserved long-distance interactions, it is able to successfully model the overall shape of novel proteins consistently, even in the absence of such information. The fact that ESM-2 is based on natural language, and does not make use of MSA may explain why it, too, does better than AF2.

This study’s principal conclusion is that orphan proteins, or TRGPs, are often predicted to have a compact 3D structure, sometimes with a novel fold, derived from their novel sequences, which are associated with the appearance of new biological functions. It will be interesting to express and purify more of these proteins, so as to determine their experimental structures, in order to find out whether some of them also have novel folds.

Because the sequences of orphan proteins lack homology information, protein structure prediction for them has recently become a hot topic^79^. The approaches and methodologies that we have implemented in this study may provide a starting point for datasets and protocols to evaluate the performance of structure prediction algorithms on sequences that lack homology to other sequences.

## Supporting information

Morphs of the 5 top models from ROSETTAFold and AF2 for the seven orphan proteins.

## Acknowledgement

The Israeli tutors and the Chinese students acknowledge the support of the YutChun-Weizmann Program that enabled this study. The study was also supported by a research grant from the Center for Scientific Excellence at the Weizmann Institute of Science. We are grateful to Dr. Shifra Ben-Dor for valuable discussions, to Prof. Robin Gasser (University of Melbourne) for providing us with the sequence of HCO_011565, to Prof. Keith Dunker (University of Indiana) for recommending the flDPnn algorithm for disorder prediction, and to Dr. Sergey Ovchinnikov (Harvard University) for valuable advice concerning the use of AF2 Colab.

