## Supplementary material for "Do Newly Born Orphan Proteins Resemble Never Born Proteins? A Study Using Three Deep Learning Algorithms": Morphs of the 5 top models from ROSETTAFold and AF2 for the seven orphan proteins.

### Slide 1
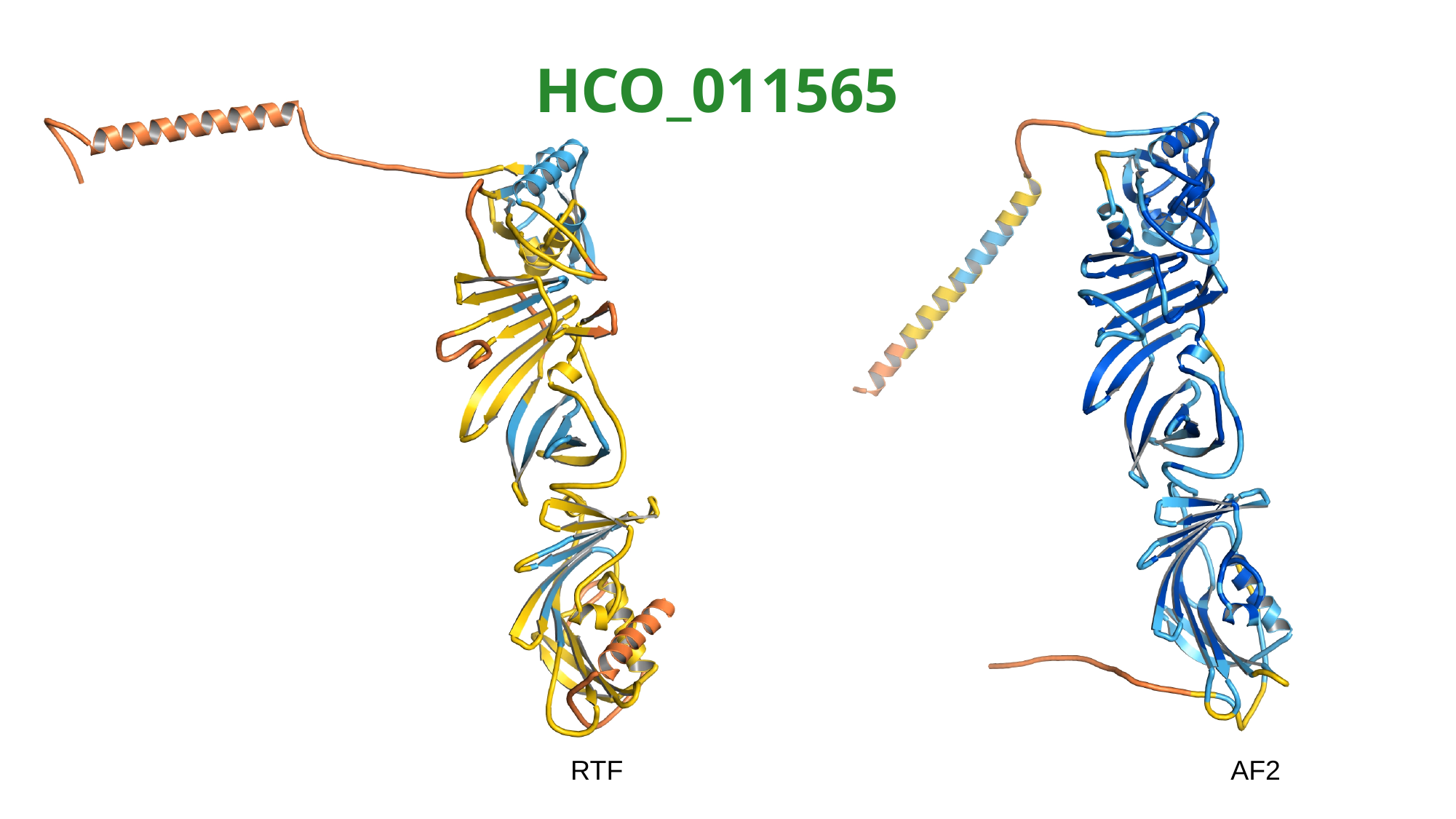

HCO_011565
RTF AF2

### Slide 2
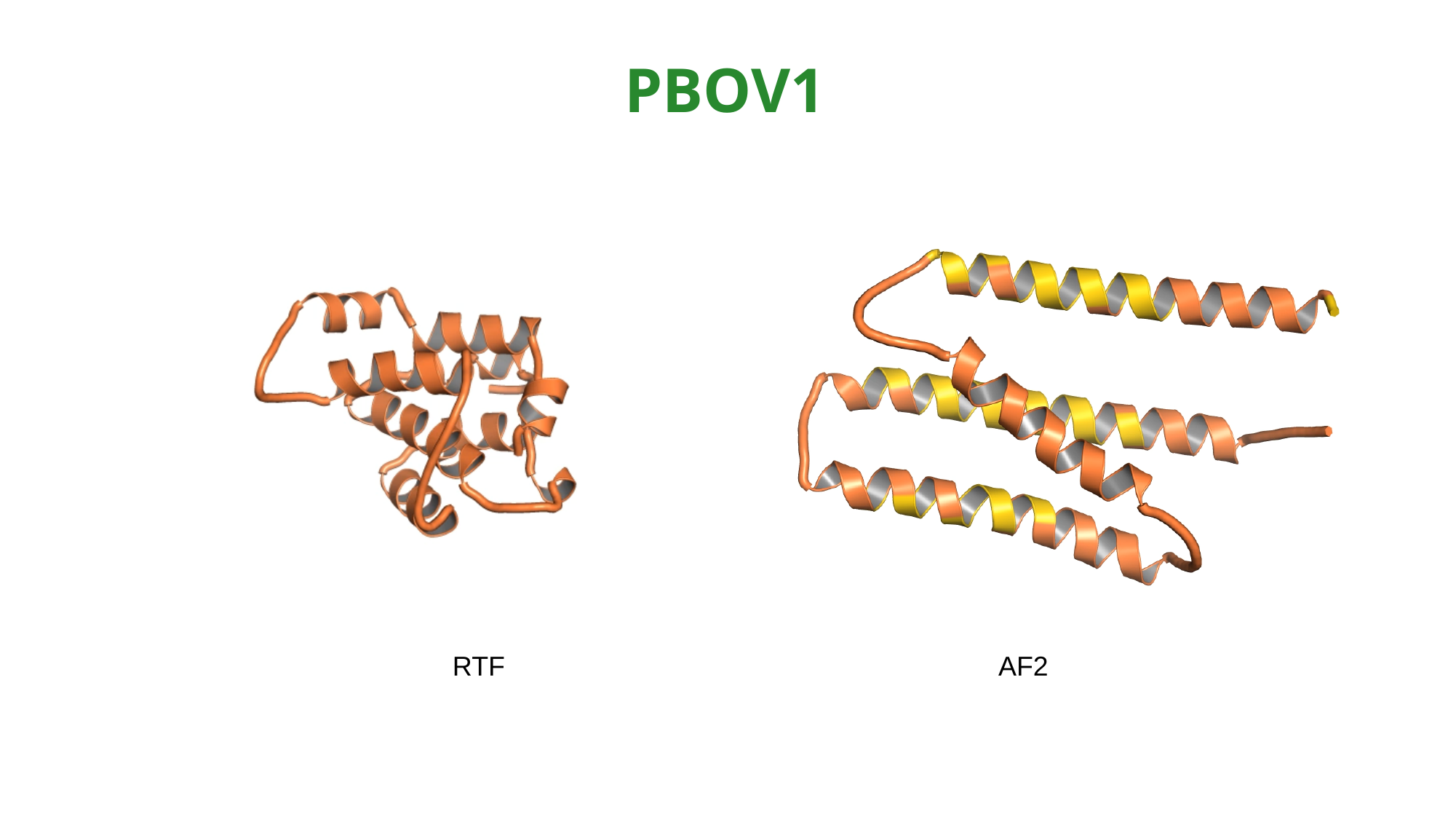

PBOV1
RTF AF2

### Slide 3
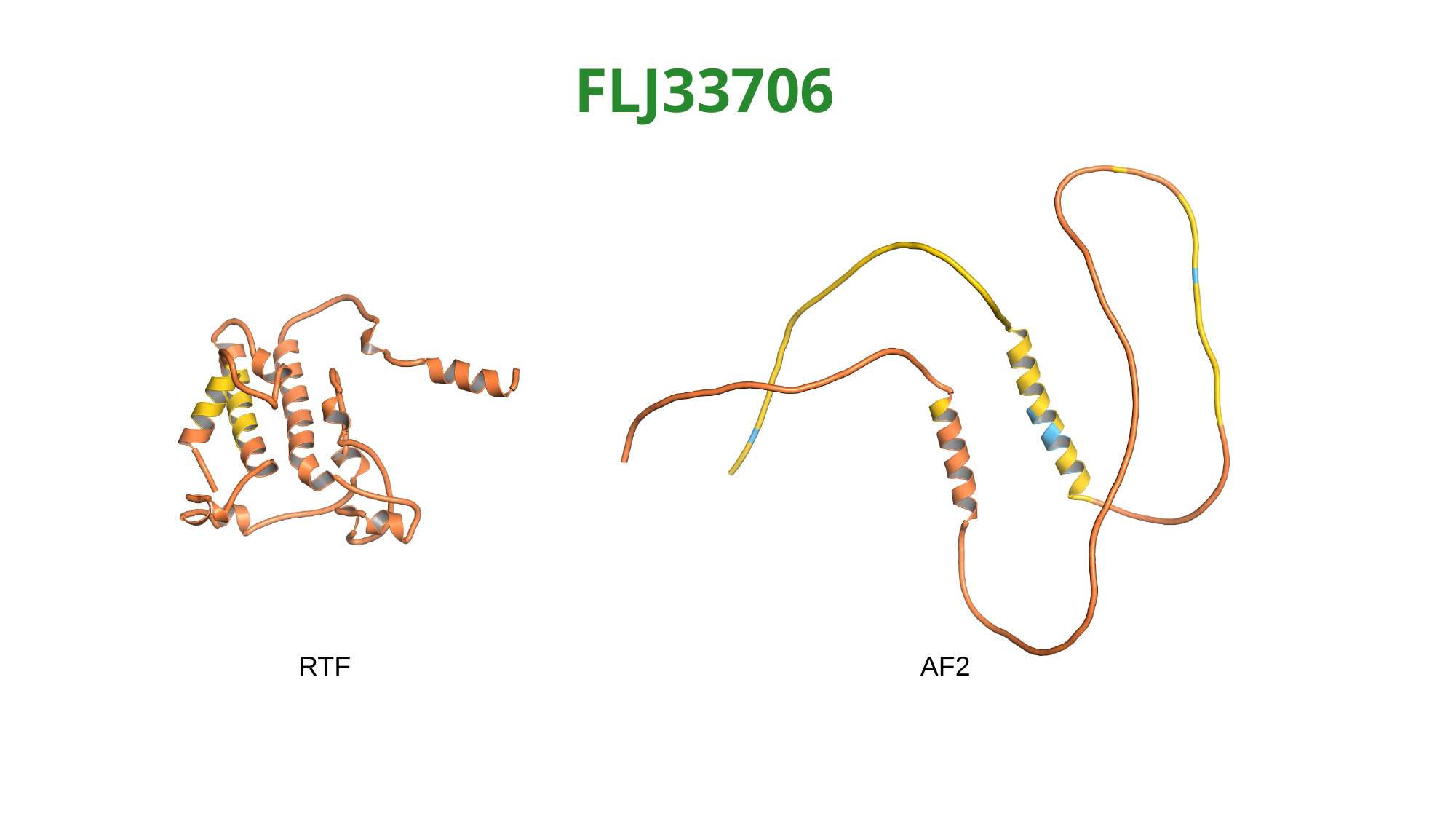

FLJ33706
 RTF AF2

### Slide 4
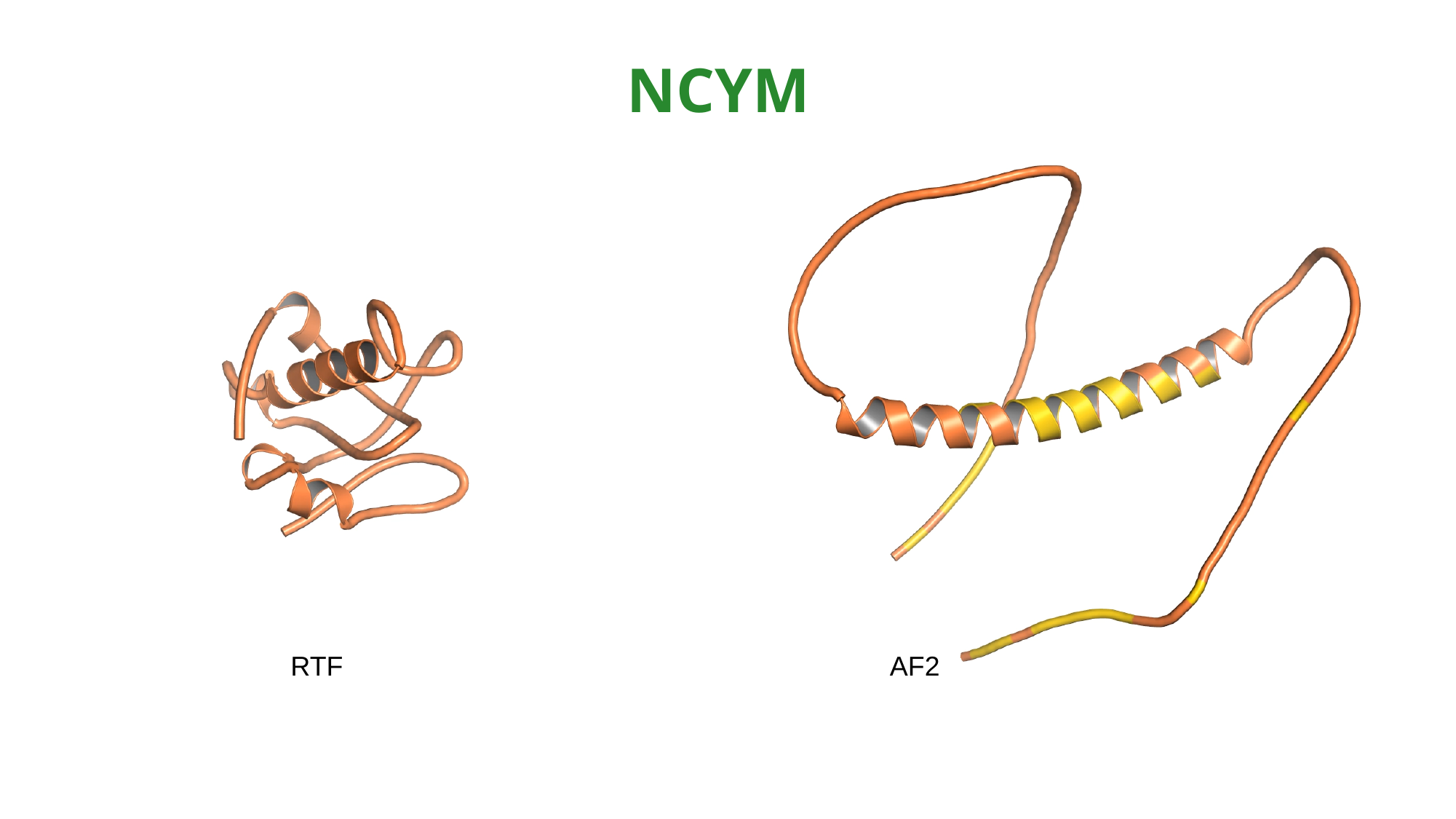

NCYM
RTF AF2

### Slide 5
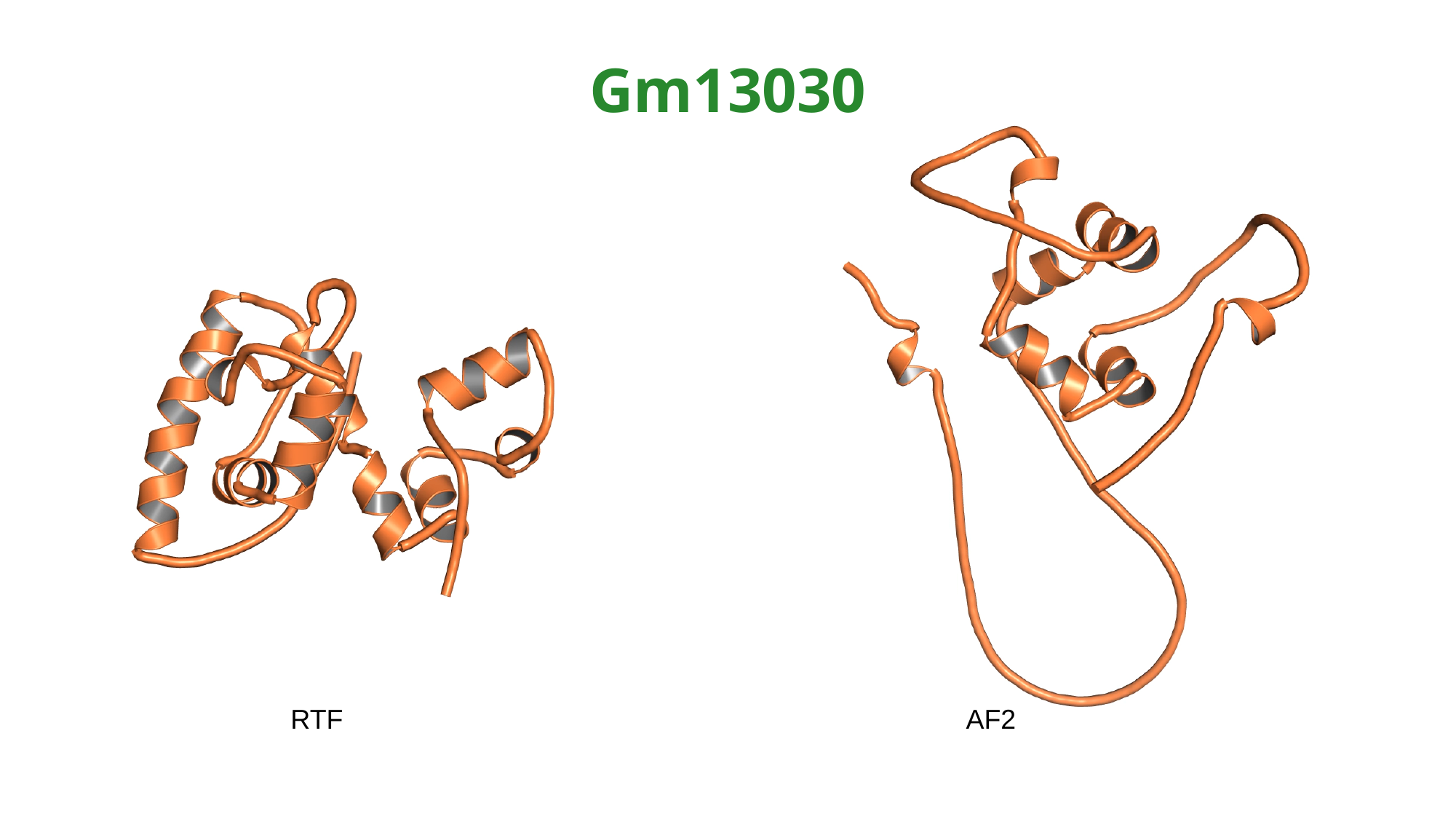

Gm13030
RTF AF2

### Slide 6
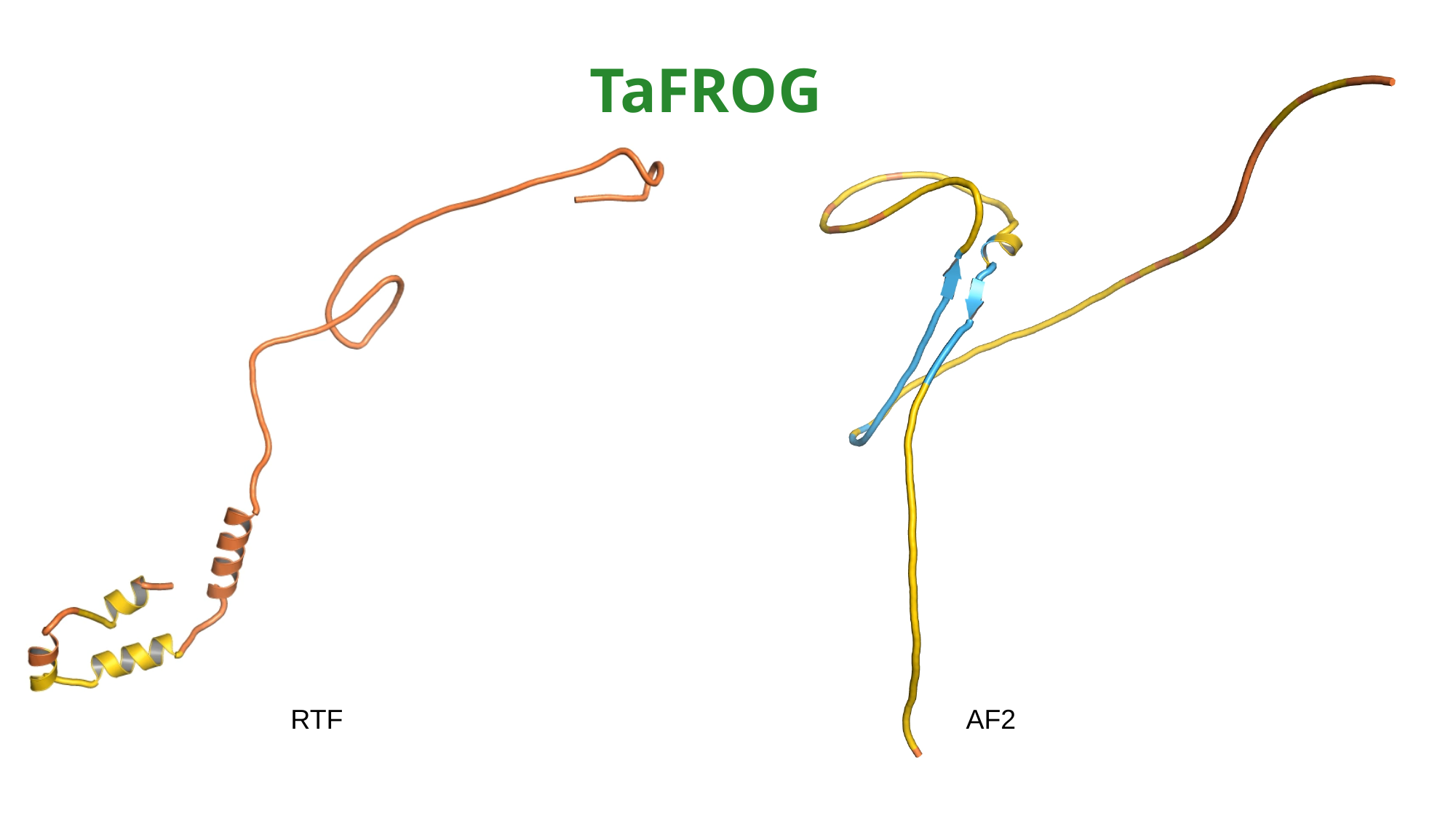

TaFROG
RTF AF2

### Slide 7
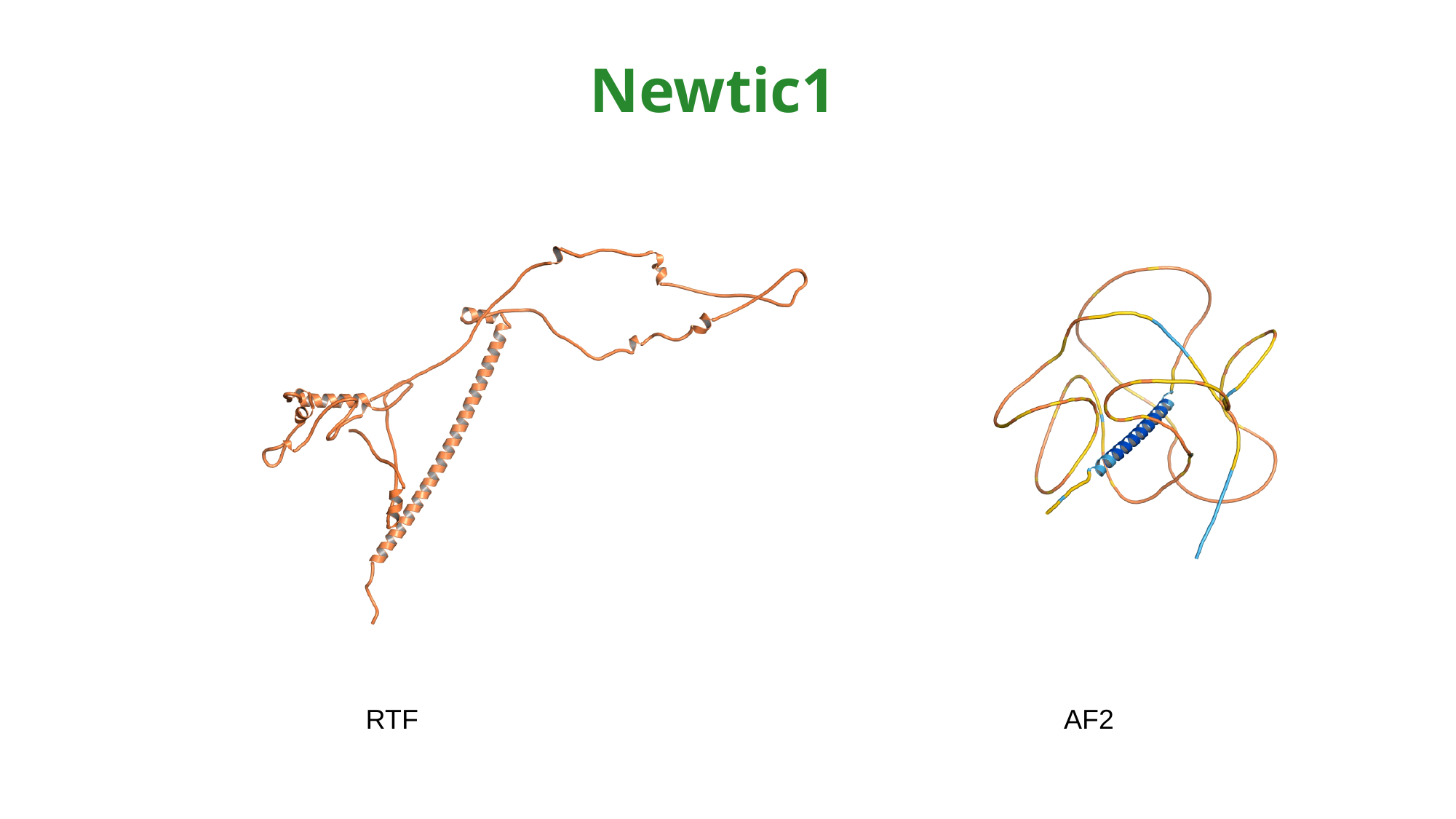

Newtic1
RTF AF2
